# Characterizing gene expression profiles of various tissue states in stony coral tissue loss disease using a feature selection algorithm

**DOI:** 10.1101/2024.11.05.622084

**Authors:** Kelsey M. Beavers, Daniela Gutierrez-Andrade, Emily W. Van Buren, Madison A. Emery, Marilyn E. Brandt, Amy Apprill, Laura D. Mydlarz

## Abstract

Stony coral tissue loss disease (SCTLD) remains a substantial threat to coral reef diversity already threatened by global climate change. Restoration efforts and effective treatment of SCTLD requires an in-depth understanding of its pathogenesis in the coral holobiont as well as mechanisms of disease resistance. Here, we present a supervised machine learning framework to describe SCTLD progression in a major reef-building coral, *Montastraea cavernosa*, and its dominant algal endosymbiont, *Cladocopium goreaui*. Utilizing support vector machine recursive feature elimination (SVM-RFE) in conjunction with differential expression analysis, we identify a subset of biologically relevant genes that exhibit the highest classification performance across three types of coral tissues collected from a natural reef environment: apparently healthy tissue on an apparently healthy colony, apparently healthy tissue on a SCTLD-affected colony, and lesion tissue on a SCTLD-affected colony. By analyzing gene expression signatures associated with these tissue health states in both the coral host and its algal endosymbiont (family Symbiodiniaceae), we describe key processes involved in SCTLD resistance and disease progression within the coral holobiont. Our findings further support evidence that SCTLD causes dysbiosis between the coral host and its Symbiodinaiceae and additionally describes the metabolic and immune shifts that occur as the holobiont transitions from a healthy to a diseased state. This supervised machine learning framework offers a novel approach to accurately assess the health states of endangered coral species and brings us closer to developing effective solutions for disease monitoring and intervention.

**AUTHOR SUMMARY:** Coral reefs are under increasing threat due to climate change, with rising ocean temperatures and disease outbreaks accelerating reef degradation. Stony coral tissue loss disease (SCTLD) has been particularly destructive, leading to widespread coral mortality across Florida’s Coral Reef and the wider Caribbean since its emergence in 2014. While the cause of SCTLD remains unknown, the rapid decline in coral reef health highlights the urgent need for innovative approaches to understanding threats to coral health. In this study, we applied a supervised machine learning approach, previously used in cancer research, to identify key genes associated with SCTLD progression in the coral *Montastraea cavernosa* and its symbiotic algae, which the coral relies on to meet its nutritional requirements. By analyzing gene expression patterns across tissues representing different health states, we find that SCTLD affects the metabolic interactions between the coral and their symbionts and causes shifts in coral immune signaling pathways, even in tissue on a SCTLD-affected colony that appears to be healthy. This study presents a novel framework for applying supervised machine learning in coral gene expression research and could lead to new methods for monitoring coral health and combatting SCTLD.

## INTRODUCTION

Coral reefs are among the most threatened ecosystems on the planet. Anthropogenic ocean warming among other stressors has triggered mass bleaching and disease outbreaks, resulting in substantial coral cover and reef biodiversity loss on nearly all the world’s tropical coral reefs [1–6]. Recovery windows between stress events have narrowed at an unprecedented rate [7]. This limits the ability of reefs to recover without intervention, highlighting the importance of coral reef management efforts as well as meaningful action on climate change [8–10]. Recent efforts have focused on the identification of coral species, individuals, symbiont genera, genetic mutations, and gene expression patterns that can withstand multiple types of stressors [11–16]. The rapid expansion of high-throughput (omics) datasets such as genomics, transcriptomics, and metabolomics and the advancement of novel statistical methods such as machine learning hold promise for the accurate identification and characterization of emerging and persistent threats to coral reef ecosystems.

One disease event that has significantly altered coral reef assemblages and functionality is stony coral tissue loss disease (SCTLD) [17–21]. First observed off the coast of Miami, Florida in 2014, SCTLD has led to significant losses of coral throughout Florida’s Coral Reef and the wider Caribbean [22–27] and is the most pervasive and virulent coral disease on record. Despite research efforts, the etiology of SCTLD remains unknown, likely due to the complexity of microbial and eukaryotic assemblages that associate with coral [1,28–30]. However, shifts in the bacterial [31–38] and viral [39,40, but see 41] consortium have been implicated in SCTLD progression, and antibiotic treatment has proven effective at halting active lesion progression [42–45]. Furthermore, histopathology has identified SCTLD tissue necrosis originating in the coral gastrodermis where the coral’s algal endosymbionts (Symbiodiniaceae) reside [46], as well as morphologic changes consistent with Symbiodiniaceae pathology in apparently healthy and SCTLD-affected corals [39], leading to the hypothesis that SCTLD may be caused by an infection of Symbiodiniaceae rather than the coral host itself.

Recent omics analyses have provided further insight into the cellular mechanisms underpinning SCTLD pathogenesis. Metabolomics on apparently healthy and SCTLD-affected corals has revealed variations in Symbiodiniaceae-derived lipid and tocopherol production in response to disease [47], supporting the case for Symbiodiniaceae involvement. Transcriptomic analyses have identified signatures of *in situ* degradation of photosynthetically dysfunctional Symbiodiniaceae [48] as well as commonly differentially expressed genes involved in innate immunity, apoptosis and extracellular matrix (ECM) structure in SCTLD-affected corals [49,50]. Interestingly, paired *ex situ* transmission and *in situ* intervention experiments showed that amoxicillin treatment led to a ‘reversal’ of many of these signaling pathways, suggesting that disease intervention provides benefits to the coral beyond removal of pathogens and opportunistic microbes [50]. Despite these advances in our understanding of SCTLD, there is still a need to identify the central processes involved in SCTLD progression to assist coral preservation and restoration projects.

Next-generation sequencing technologies have revolutionized disease research, allowing scientists to measure expression-level changes of thousands of genes in response to experimental or natural infection [51]. However, gene expression datasets are high-dimensional in nature, and the differentially expressed genes (DEGs) produced often contain redundant and biologically irrelevant data [52]. Integrating supervised machine learning (ML), defined as the process of learning from labelled examples to predict or classify an outcome of interest [53], into DEG analyses offers a promising solution to address the ‘curse of dimensionality’ [54] prevalent in large omics datasets. One widely adopted supervised ML algorithm for feature selection is Support Vector Machine Recursive Feature Elimination (SVM-RFE). SVM-RFE identifies the features (genes) in a dataset that optimally define the class (phenotype) data [55] and has many applications in gene expression research. For example, SVM-RFE has been used to predict drought-resistant genes in *Arabidopsis thaliana* [56], identify genes for accurate cancer classification [55], and diagnose Alzheimer’s disease in humans [57]. Broadly, feature selection using SVM-RFE has been shown to be successful at isolating a subset of nonredundant and biologically relevant genes from a larger dataset, enabling the construction of robust models capable of accurately assigning data points to predefined classes.

Here, we implement a novel supervised ML approach aimed at characterizing the gene expression associated with various tissue health states in a major reef-building coral *Montastraea cavernosa,* and its dominant algal endosymbiont, *Cladocopium goreaui*. Through the use of the “sigFeature” feature selection algorithm [52], we implement a simultaneous combination of SVM-RFE and differential expression analysis to identify genes that are both biologically relevant and have the highest discriminatory power within three types of coral tissue collected from a natural reef environment: apparently healthy tissue on an apparently healthy colony, apparently healthy tissue on a SCTLD-affected colony, and lesion tissue on a SCTLD-affected colony. This approach allows us to pinpoint the most relevant genes associated with different states of SCTLD progression, thereby enhancing our understanding of its pathogenesis and facilitating the development of effective treatment and prevention strategies.

## RESULTS

### Transcriptome assembly and annotation

Sequencing of 22 coral tissue samples resulted in a total of 1.035 billion raw reads with an average of 47 million raw reads per sample. *De novo* transcriptome assembly of the cleaned and quality-filtered *M. cavernosa* reads resulted in an assembly of 73,047 contigs with a N50 size of 16,467 bp and filtering of the *C. goreaui de novo* transcriptome published previously [58] resulted in an assembly of 48,013 contigs with a N50 size of 13,469 bp (Table 1). Of those, 33,614 *M. cavernosa* and 26,245 *C. goreaui* contigs were annotated with an annotation evalue < 1.0e^−6^.

**Table 1:**
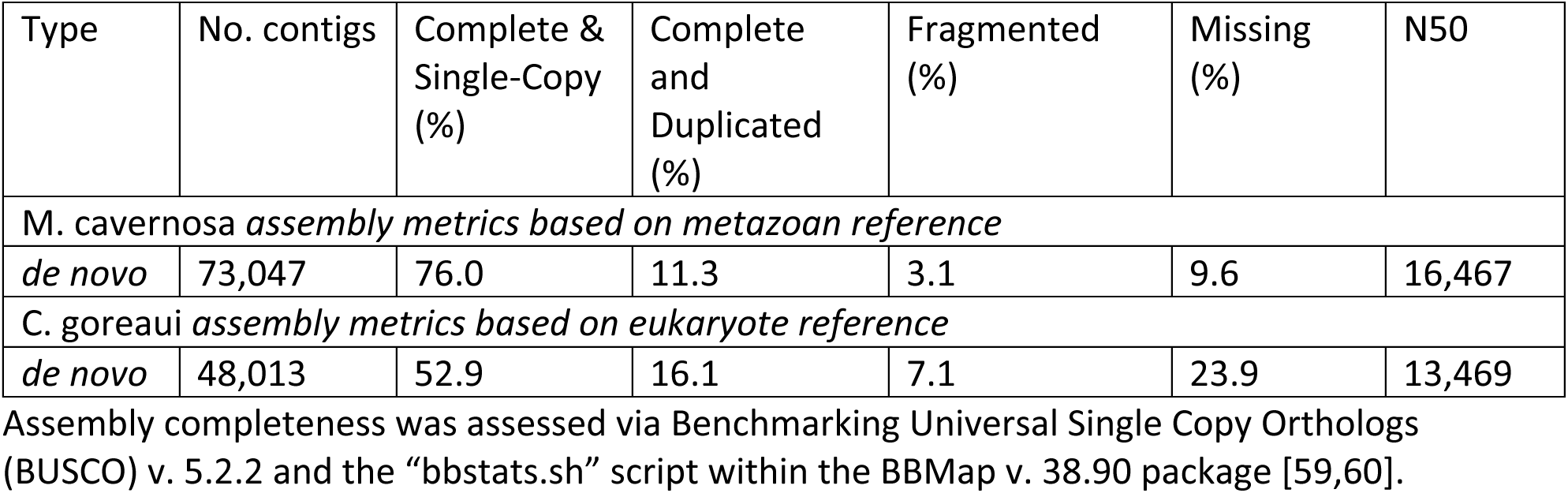
Reference transcriptome assembly metrics.

### Isolation and quantification of holobiont reads

A total of 692 million reads were assigned to *M. cavernosa* and 139 million to *C. goreaui*. An average of 31.5 million reads and 6.34 million reads were assigned to *M. cavernosa* and *C. goreaui* per sample, respectively. Mapping of *M. cavernosa* and *C. goreaui* reads to their respective refence transcriptome resulted in average mapping rates of 92.2% and 91.5%, respectively (Table S1). Following transcript quantification, a total of 19,039 and 11,289 length-normalized transcripts with an annotation evalue < 1.0e^−6^ were expressed in *M. cavernosa* and *C. goreaui*, respectively. Filtering out genes with an average rlog expression < 10 resulted in 17,229 and 2,224 genes with significant levels of expression in *M. cavernosa* and *C. goreaui*, respectively.

### Feature selection on holobiont gene expression

The 17,229 annotated genes with significant expression in *M. cavernosa* were used to produce the feature ranked lists from the HH, HD, and LD coral gene expression datasets (Tables S2-7). Of the *M. cavernosa* features, 1,562 HH, 5,286 HD, and 5,329 LD had significant differential expression relative to the other two tissue health states (*t*-statistic *P*-value < 0.05). External stratified *k*-fold cross-validation (*k =* 10) showed high classification performance of each tissue health state’s top 400 coral features: within the HH dataset and LD datasets, 0% average misclassification was achieved with the top 390 and 130 features, respectively, and within the HD dataset, 12% average misclassification was achieved with the top 370 features (Fig 1E).

**Fig 1.**
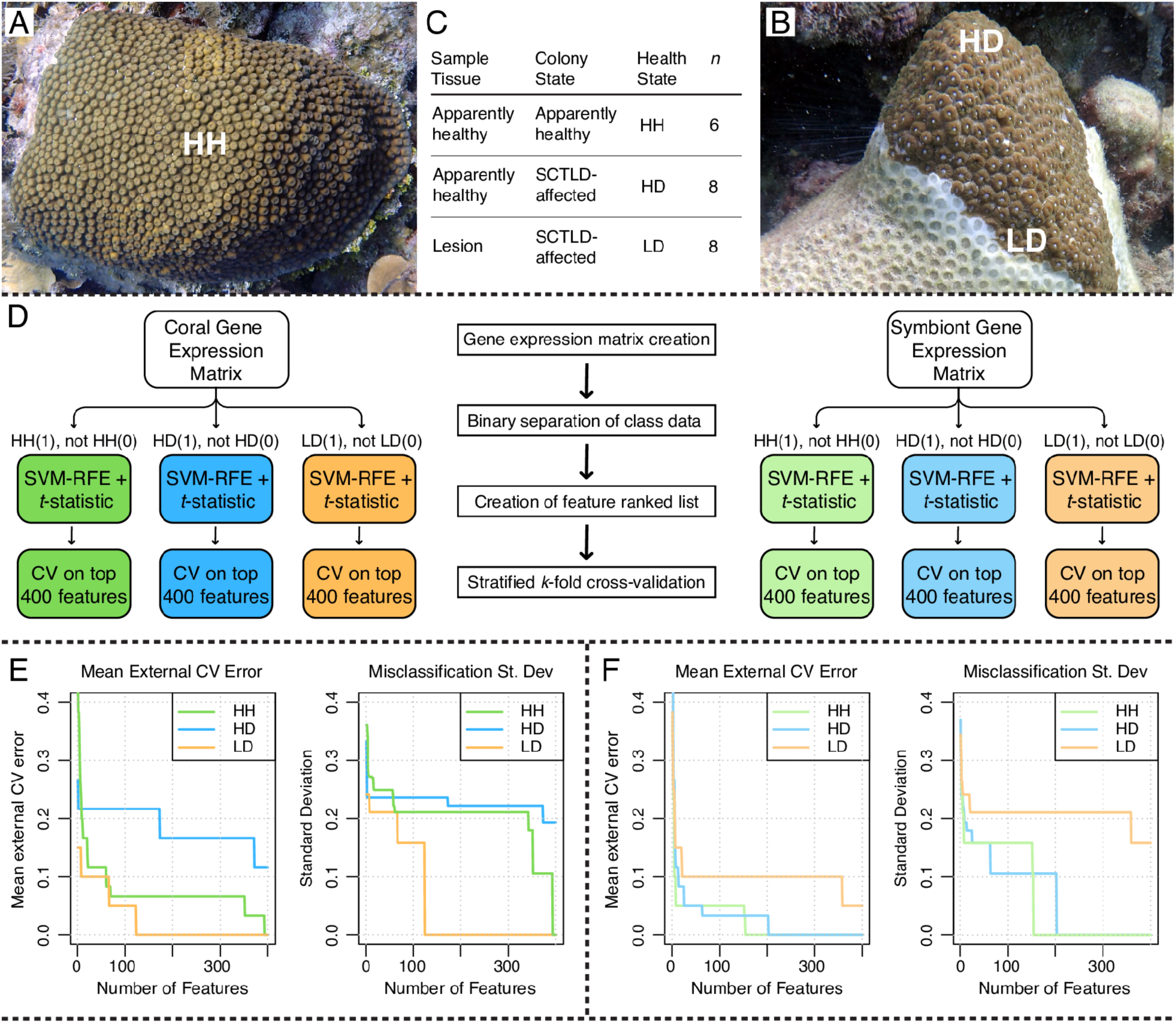
Experimental design and analysis workflow. (A-B) Photographs showing the three tissue health states collected: apparently healthy tissue on an apparently healthy colony (HH), apparently healthy tissue on a diseased colony (HD), and lesion tissue on a diseased colony (LD). (C) Sampling data. (D) Feature selection pipeline using a combination of SVM-RFE and differential expression *t-*statistic using the sigFeature R package [52]. (E) External stratified *k-*fold cross-validation results on the top 400 *M. cavernosa* features from each tissue health state. (F) External stratified *k-*fold cross-validation results on the top 400 *C. goreaui* features from each tissue health state (SVM-RFE support vector machine recursive feature elimination, CV cross-validation). (A), (B) Amy Apprill

The 2,158 annotated genes with significant expression in *C. goreaui* were used to produce the feature ranked lists from the HH, HD and LD symbiont gene expression datasets. Of the *C. goreaui* features, 152 HH, 114 HD, and 291 LD had a significant differential expression relative to the other two tissue health states (*t-*statistic *P*-value < 0.05). External stratified *k*-fold cross-validation (*k =* 10) showed high classification performance of each tissue health state’s top 400 symbiont features: within the HH and HD datasets, 0% average misclassification was achieved with the top 160 and 210 features respectively, and within the LD dataset, 5% average misclassification was achieved with the top 360 features (Fig 1F).

### Functional enrichment of top features in M. cavernosa

Of the top 500 *M. cavernosa* HH features, 369 were upregulated and 130 were downregulated relative to both HD and LD tissue. Functional enrichment of those genes showed that apparently healthy tissue on an apparently healthy *M. cavernosa* colony is characterized by increased unsaturated fatty acid biosynthesis, collagen formation, and actin binding as well as decreased nitric oxide biosynthesis, glycogen biosynthesis, and amino acid catabolism (Fig 2). Of the top 500 *M. cavernosa* HD features, 202 were upregulated and 297 were downregulated relative to both HH and LD tissue. Functional enrichment of those genes showed that apparently healthy tissue on a SCTLD-affected *M. cavernosa* colony is characterized by increased translation, amide biosynthesis, and aminoacyl-tRNA biosynthesis as well as decreased cilium movement and assembly and mitochondrion organization (Fig 2). Of the top 500 *M. cavernosa* LD features, 229 were upregulated and 271 were downregulated relative to both HH and HD tissue. Functional enrichment of those genes showed that lesion tissue on a SCTLD-affected *M. cavernosa* colony is characterized by increased innate immunity through C-type lectin receptor pattern recognition, NF-kappaB (NF-κB) signaling, and autophagosome maturation as well as decreased collagen chain trimerization, chloride transmembrane transport, and unsaturated fatty acid biosynthesis (Fig 2).

**Fig 2.**
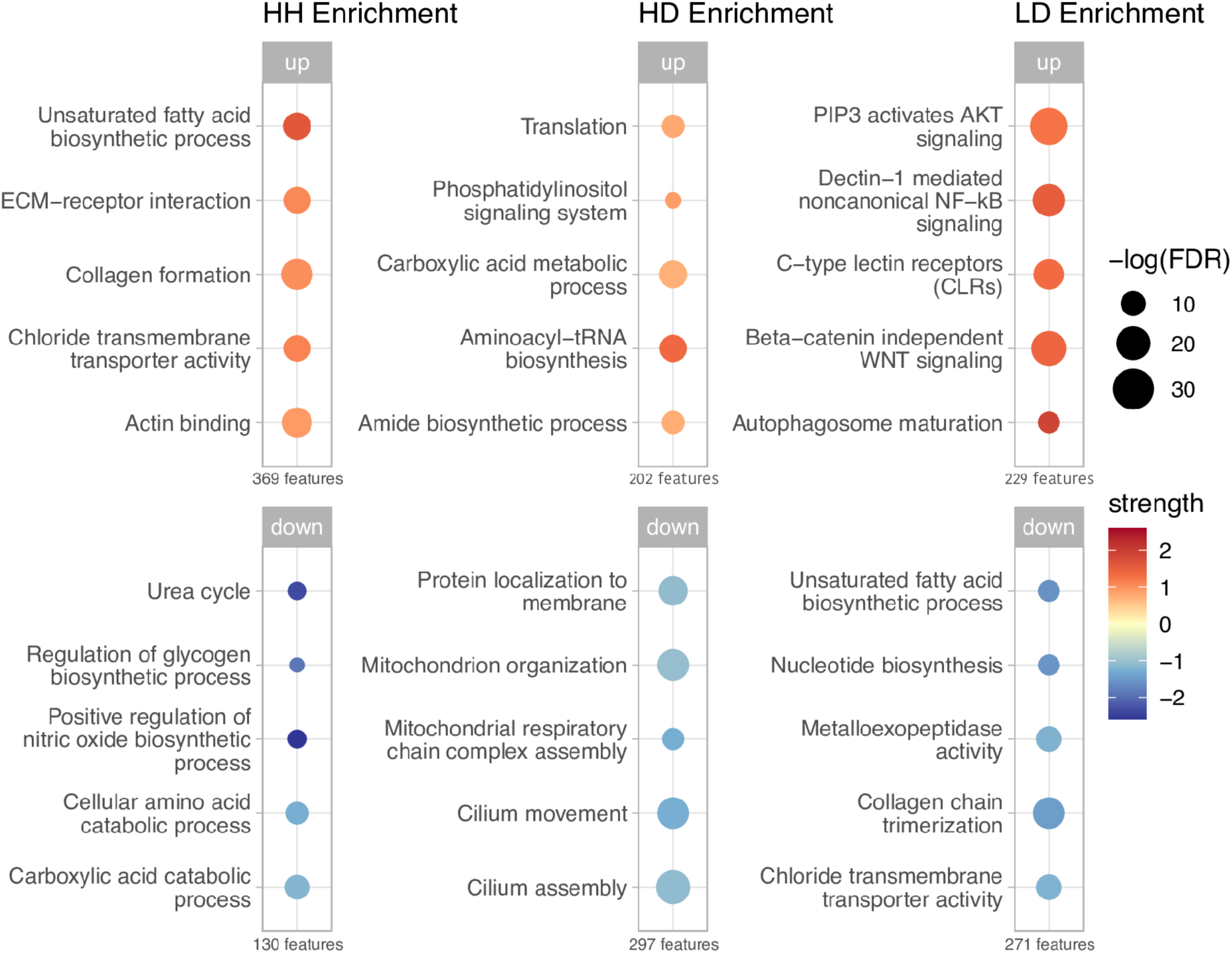
Functional enrichment of up- and downregulated genes within the top 500 *M. cavernosa* features from each tissue health state. The number of genes (features) in each list of up- and downregulated genes from each tissue health state are shown beneath each bubble plot. Upregulated enrichments are shown on the top panel and downregulated enrichments are shown on the bottom panel. Size of bubble represents the –log transformed false discovery rate for the enrichment, corrected for multiple testing using Benjamini–Hochberg procedure. Color represents the strength of the enrichment: the ratio between the number of proteins in the network that are annotated and the number of proteins that we expect to be annotated with this term in a random network of the same size (Log10(observed/expected)). Strength values from the downregulated enrichments were multiplied by −1 for visualization purposes.

Of the top 500 *C. goreaui* HH features, 259 were upregulated and 225 were downregulated relative to both HD and LD tissue. Functional enrichment of those genes showed that *C. goreaui* from apparently healthy tissue on an apparently healthy *M. cavernosa* colony is characterized by increased long-chain fatty acid biosynthesis, heat shock responses, and sphingolipid metabolism as well as decreased pyruvate metabolism and phenylpropanoid biosynthesis (Fig 3). Of the top 500 *C. goreaui* HD features, 224 were upregulated and 221 were downregulated relative to both HH and LD tissue. Functional enrichment of those genes showed that *C. goreaui* from apparently healthy tissue on a SCTLD-affected colony is characterized by increased organonitrogen and organophosphate biosynthesis as well as decreased regulation of the mitotic cell cycle and iron uptake and transport (Fig 3). Of the top 500 *C. goreaui* LD features, 224 were upregulated and 276 were downregulated relative to both HH and HD tissue. Functional enrichment of those genes showed that *C. goreaui* from lesion tissue on a SCTLD-affected colony is characterized by increased thiamine metabolism and proteasomal protein catabolism as well as decreased inorganic anion transmembrane transport, chloride channel activity, and signaling by nuclear receptors (Fig 3).

**Fig 3.**
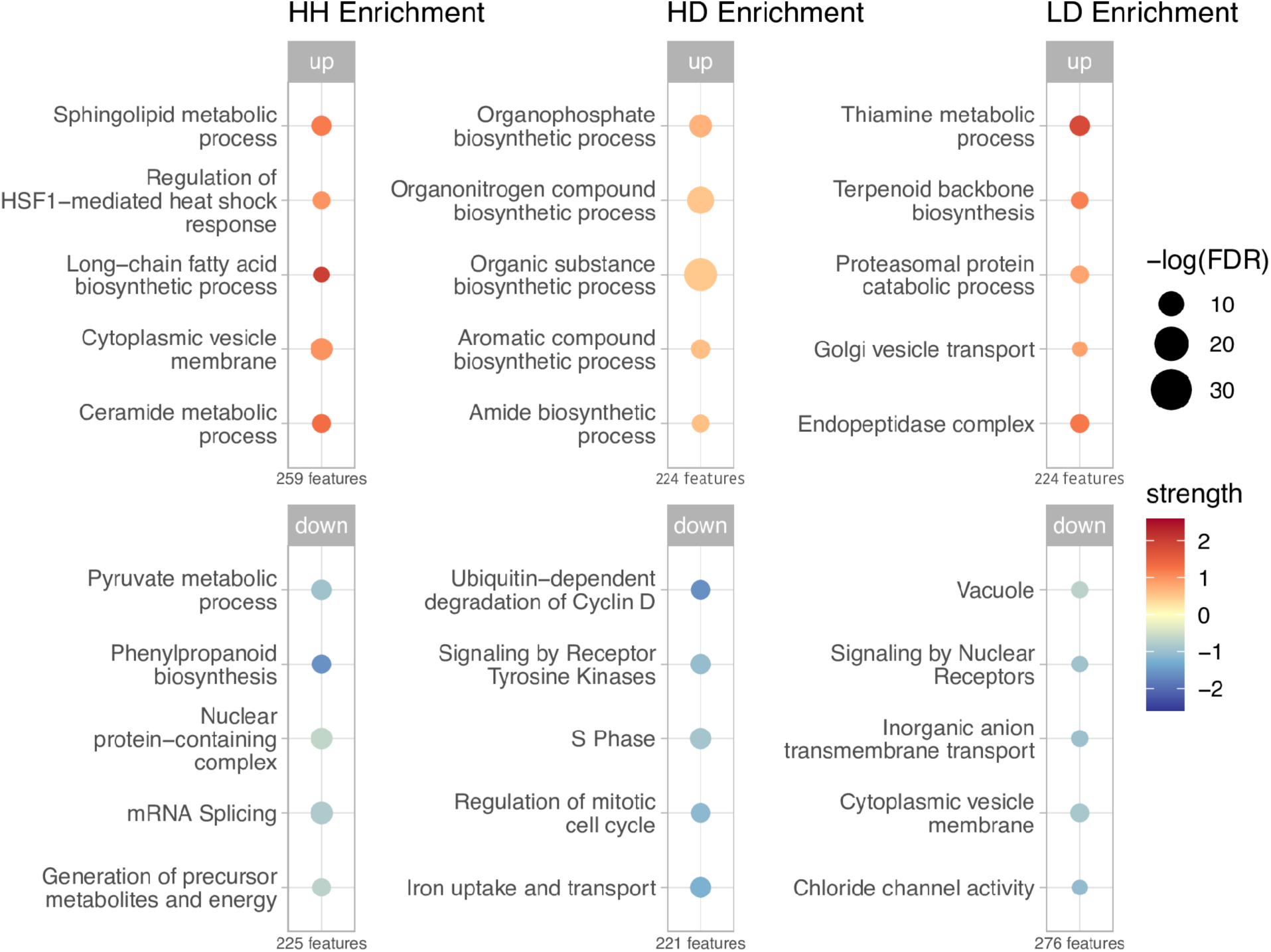
Functional enrichment of up- and downregulated genes within the top 500 *C. goreaui* features from each tissue health state. The number of genes (features) in each list of up- and downregulated genes from each tissue health state are shown beneath each bubble plot. Upregulated enrichments are shown on the top panel and downregulated enrichments are shown on the bottom panel. Size of bubble represents the –log transformed false discovery rate for the enrichment, corrected for multiple testing using Benjamini–Hochberg procedure. Color represents the strength of the enrichment: the ratio between the number of proteins in the network that are annotated and the number of proteins that we expect to be annotated with this term in a random network of the same size (Log10(observed/expected)). Strength values for the downregulated enrichments were multiplied by −-1 for visualization purposes.

### Expression analysis of potential tissue health state biomarkers

Of the top 15 HH features in *M. cavernosa*, 13 (86.7%) were upregulated and two (13.3%) were downregulated relative to both HD and LD tissue (Fig 4A). The top feature in HH corals was the upregulated Transmembrane protein 145 (*Tmem145*), an integral membrane component associated with transforming growth factor beta (TGFβ) signaling (Table S14) [61]. Other notable genes within the top 15 HH features include: Prokineticin receptor 1 (*PROKR1*), Inhibin beta B chain (*INHBB*), Baculoviral IAP repeat-containing protein 1 (*NAIP*) and Deleted in malignant brain tumors 1 protein (*Dmbt1*) (Fig 4). Of the top 15 HD features in *M. cavernosa*, five (33.3%) were upregulated and ten (66.7%) were downregulated relative to both HH and LD tissue (Fig 5A). The top feature in HD corals was the upregulated Dicarboxylate carrier SLC25A8 (*Ucp2*), a mitochondrial uncoupling protein that primarily functions in decreasing mitochondrial membrane potential and reactive oxygen species (ROS) production (Table S15) [61]. Other notable genes within the top 15 HD features include: Ras-related protein Rab-5A (*Rab5a*), Cilia- and flagella-associated protein 298 (*cfap298),* Ciliary microtubule inner protein 2B (*fam166b*), and Chaperone protein DnaJ (*dnaJ*) (Fig 5). Of the top 15 LD features in *M. cavernosa*, 14 (93.3%) were upregulated and one (6.7%) was downregulated relative to both HH and HD tissue (Fig 6A). The top feature in LD corals was the downregulated Tropomyosin-1 (*TPM1*), a non-muscle tropomyosin isoform involved in the control and regulation of the cell’s cytoskeleton (Table S16) [61]. Other notable genes within the top 15 LD features include: two Collagen alpha-1(II) chain homologs (*col2a1 [a]* and *col2a1 [b]*), two Collagen alpha-2(I) chain homologs (*col1a2 [a]* and *col1a2 [b]*), GFP-like fluorescent chromoprotein cFP484 (*GFPL*), and Superoxide dismutase [Cu-Zn] 1 (*sodA*) (Fig 6).

**Fig 4.**
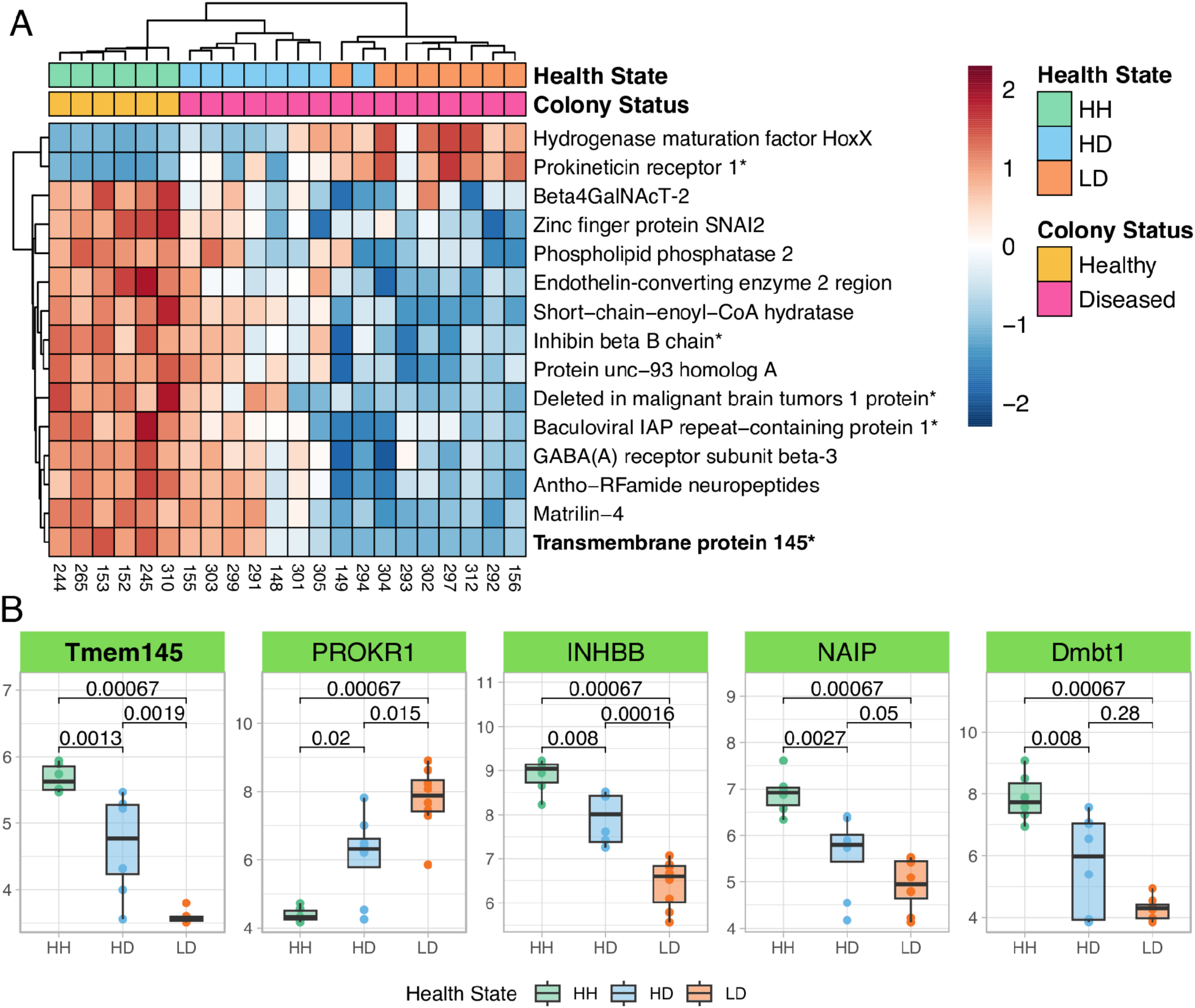
Top features in HH *M. cavernosa*. (A) Relative expression heatmap of the top 15 HH features from *M. cavernosa*. Red boxes signify elevated expression relative to the row mean, and blue boxes signify lowered expression relative to the row mean. The top feature is shown in bold, and genes plotted in (B) end in an asterisk. (B) Boxplots showing the rlog transformed expression of five selected features from (A), organized by tissue health state. *P*-values represent two-sample Wilcoxon test results. Color of boxplots correspond to tissue health state. Boxplot elements: center line, median; box limits, upper and lower quartiles; whiskers, 1.5x interquartile range; points beyond whiskers, outliers.

**Fig 5.**
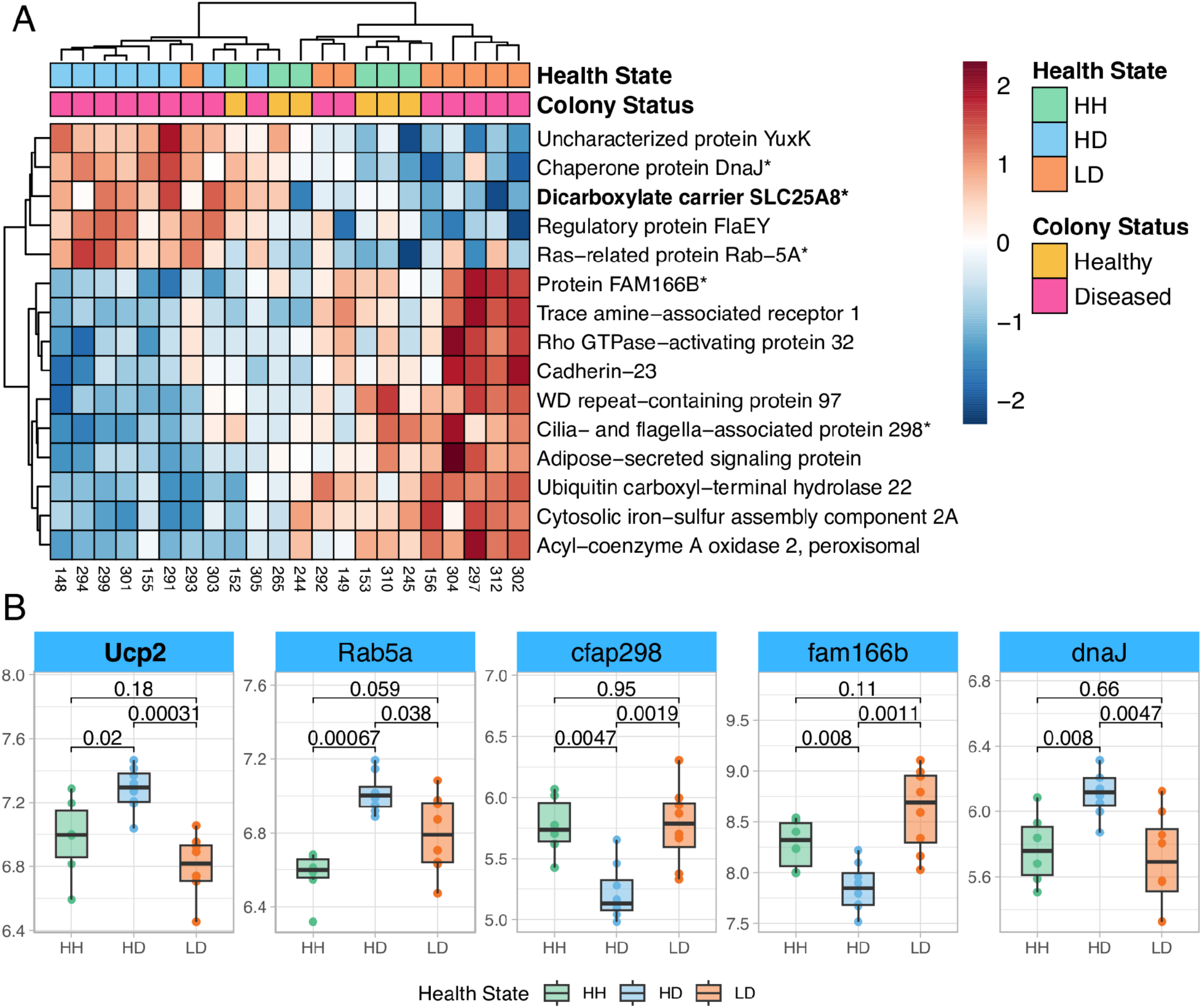
Top features in HD *M. cavernosa*. (A) Relative expression heatmap of the top 15 HD features from *M. cavernosa*. Red boxes signify elevated expression relative to the row mean, and blue boxes signify lowered expression relative to the row mean. The top feature is shown in bold, and genes plotted in (B) end in an asterisk. (B) Boxplots showing the rlog transformed expression of five selected features from (A), organized by tissue health state. *P*-values represent two-sample Wilcoxon test results. Color of boxplots correspond to tissue health state. Boxplot elements: center line, median; box limits, upper and lower quartiles; whiskers, 1.5x interquartile range; points beyond whiskers, outliers.

**Fig 6.**
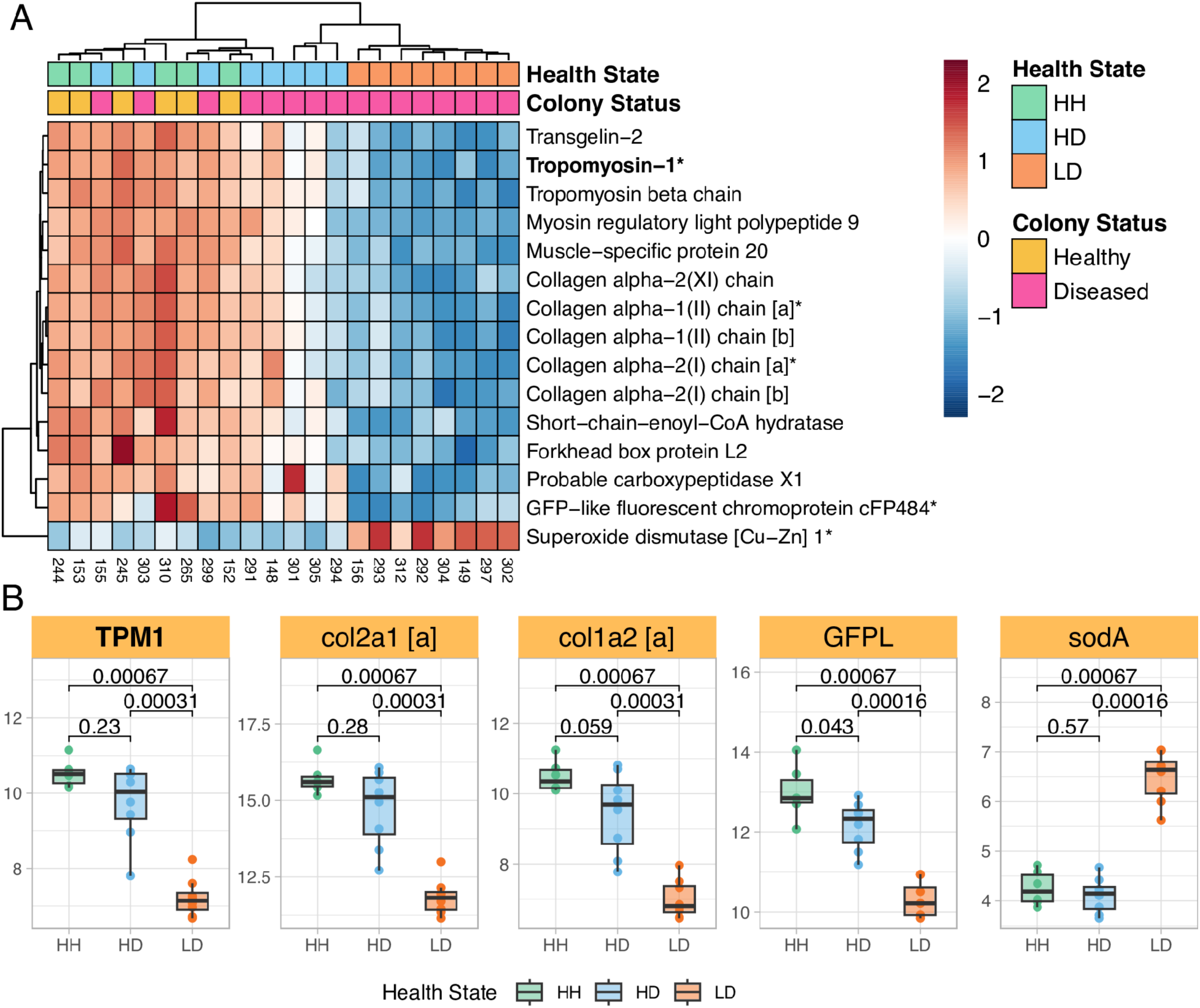
Top features in LD *M. cavernosa*. (A) Relative expression heatmap of the top 15 LD features from *M. cavernosa*. Red boxes signify elevated expression relative to the row mean, and blue boxes signify lowered expression relative to the row mean. The top feature is shown in bold, and genes plotted in (B) end in an asterisk. (B) Boxplots showing the rlog transformed expression of five selected features from (A), organized by tissue health state. *P*-values represent two-sample Wilcoxon test results. Color of boxplots correspond to tissue health state. Boxplot elements: center line, median; box limits, upper and lower quartiles; whiskers, 1.5x interquartile range; points beyond whiskers, outliers.

Of the top 15 HH features in *C. goreaui*, six (40%) were upregulated and nine (60%) were downregulated relative to both HD and LD tissue (Fig S2). The top feature in HH *C. goreaui* symbionts was the upregulated Protein NO VEIN, which contains a histidine kinase/HSP90-like ATPase superfamily domain found in several ATP-binding proteins (Table S17) [61]. Other notable genes within the top 15 HH features include Inner membrane ALBINO3-like protein 2, chloroplastic (*ALB3.2*) and Cytochrome b5 (*Cyt-b5*) (Fig 7). Of the top 15 HD features in *C. goreaui*, seven (46.7%) were upregulated and eight (53.3%) were downregulated relative to both HH and LD tissue (Fig S3). The top feature in HD *C. goreaui* symbionts was the upregulated Aspartate ammonia-lyase (*aspA*), which converts L-aspartate to fumarate and ammonia during the tricarboxylic acid cycle (Table S18) [61]. Other notable genes within the top 15 HD features include Serine/threonine-protein kinase STY17 (*STY17*) and Soluble starch synthase 1, chloroplastic/amyloplastic (*SS1*) (Fig 7). Of the top 15 LD features in *C. goreaui*, 13 (86.7%) were upregulated and two (13.3%) were downregulated relative to both HH and HD tissue (Fig S4). The top feature in LD *C. goreaui* symbionts was the downregulated High affinity nitrate transporter 2.5 (*NRT2.5*), which is involved in the transmembrane transport of nitrogen (Table S19) [61]. Other notable genes within the top 15 LD features include Fucoxanthin-chlorophyll a-c binding protein E, chloroplastic (*FCPE*) and glycosyltransferase-like KOBITO 1 (*ELD1*) (Fig 7).

**Fig 7.**
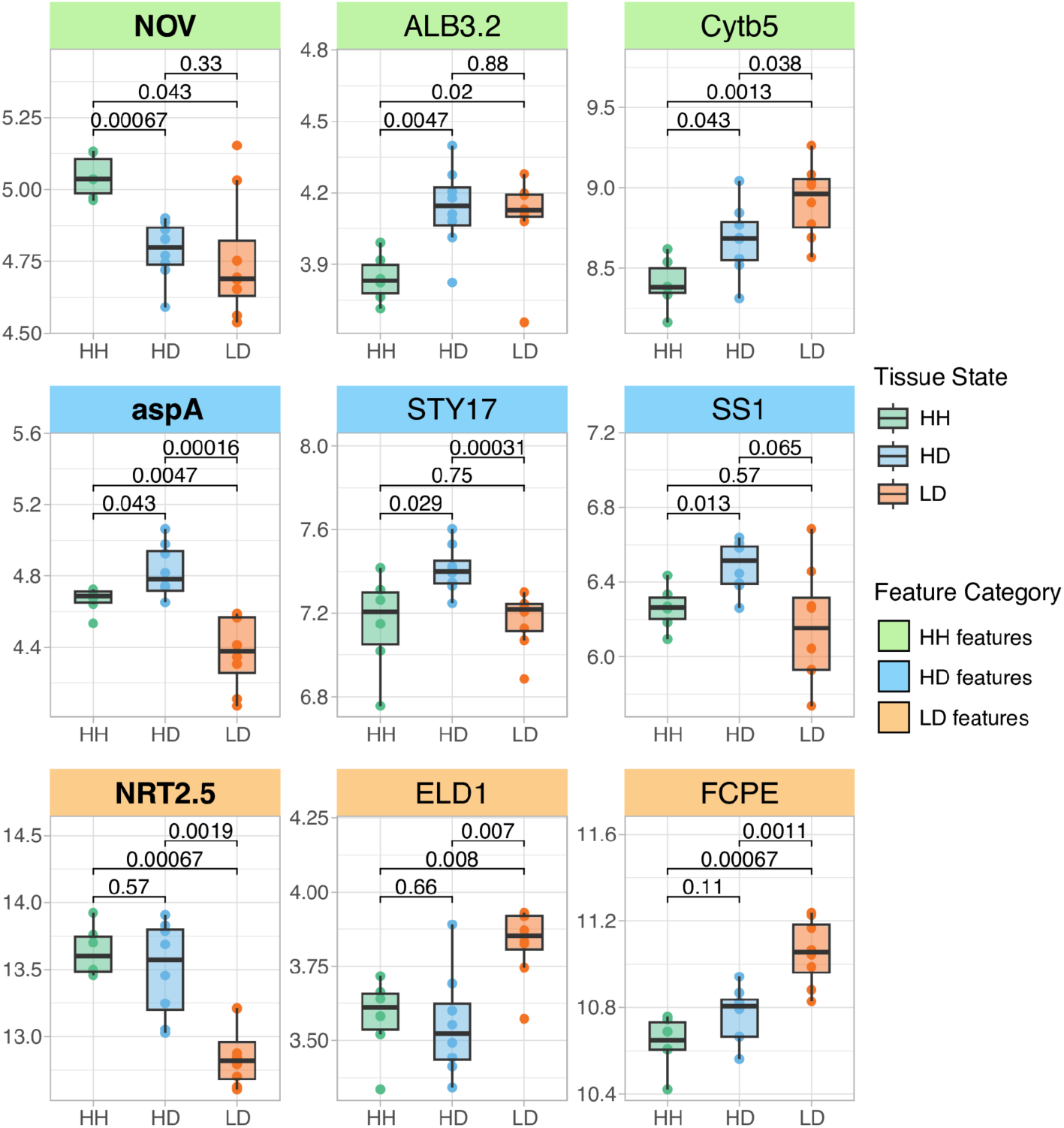
Selected top features in *C. goreaui*. Boxplots show the rlog transformed expression of three selected features from each tissue health state in *C. goreaui.* HH features are shown in the top panel, HD features in the middle panel, and LD features in the bottom panel. The top feature from each tissue health state is shown in the first column in bold. *P-*values represent two-sample Wilcoxon test results. Color of boxplots correspond to tissue health state. Boxplot elements: center line, median; box limits, upper and lower quartiles; whiskers, 1.5x interquartile range; points beyond whiskers, outliers.

## DISCUSSION

Supervised machine learning (ML) is a powerful tool that uses known characteristics to detect meaningful patterns within large, unstructured, and complex datasets [62]. Here, we use a supervised ML feature selection algorithm to identify expression patterns that best discriminate between three distinct SCTLD health states in a principal reef-building coral, *Montastraea cavernosa*, and its dominant algal endosymbiont, *Cladocopium goreaui*. By integrating differential expression analysis with support vector machine recursive feature elimination (SVM-RFE), we produce a ranked list of genes from both *M. cavernosa* and *C. goreaui* that characterize the molecular signatures associated with apparently healthy tissue on apparently healthy colonies (HH), apparently healthy tissue on SCTLD-affected colonies (HD), and lesion tissue on a SCTLD-affected colonies (LD). Our analysis reveals distinct gene expression profiles across all three health states, as evidenced by the high classification power of each dataset’s top features (Fig 8), and sheds light on the processes involved in SCTLD pathogenesis.

**Fig 8.**
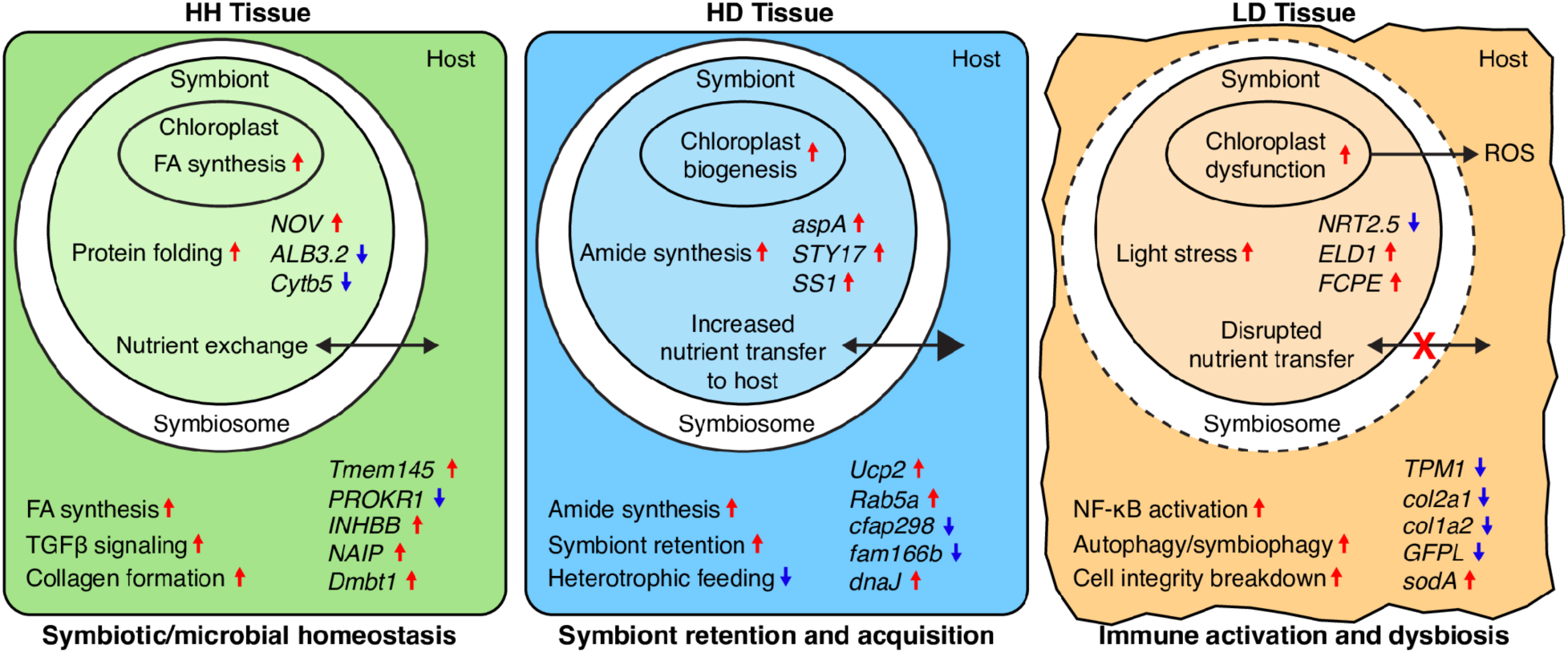
Characterization of SCTLD tissue health states in *M. cavernosa* and *C. goreaui*. Processes are shown in regular font and selected features are shown in italic font. Processes are inferred from both functional enrichment of top 500 features and analysis of selected features (genes listed). Upregulated processes/features are denoted with a red arrow and downregulated processes/features are denoted with a blue arrow. (FA, fatty acid; ROS, reactive oxygen species).

### HH Tissue is Characterized by High Fatty Acid Biosynthesis and Symbiotic Homeostasis

Metabolism within apparently healthy tissue on an apparently healthy colony is broadly characterized by high levels of fatty acid biosynthesis in both *M. cavernosa* and *C. goreaui*. Fatty acid biosynthesis is critically important for various biological processes, such as maintaining membrane structure and function, energy storage and metabolism, and cell signaling [63]. Symbiodiniaceae have been shown to supply their coral hosts with myriad saturated and polyunsaturated fatty acids [64], and previous work has found that Symbiodiniaceae-derived lipid content is higher in apparently healthy *M. cavernosa* corals as opposed to SCTLD-affected corals [47]. The enrichment of fatty acid biosynthesis in both the coral host and its algal symbiont in apparently healthy tissue further emphasizes the importance of this metabolic pathway in maintaining overall health and symbiotic homeostasis.

Within the top *M. cavernosa* HH features, we found high expression of genes involved in TGFβ signaling (*Tmem145* and *INHBB*). The TGFβ signaling pathway is conserved throughout metazoan evolution and functions in a variety of processes such as development, cellular homeostasis, and immune regulation [65]. In symbiotic cnidarians, TGFβ signaling is known to suppress host immunity to establish and maintain symbiosis with Symbiodiniaceae [66–68]. Notably, all transcriptomic studies of SCTLD to date have identified higher expression of TGFβ signaling genes in apparently healthy corals compared to their SCTLD-affected counterparts [48–50]. High constitutive levels of TGFβ signaling, therefore, may constitute an important mechanism of SCTLD resistance through the maintenance of symbiosis and prevention of a detrimental inflammatory response.

Indeed, another upregulated gene within the top *M. cavernosa* HH features includes *Dmbt1*, known to be involved in maintaining intestinal epithelia mucosal homeostasis by preventing bacterial invasion and suppressing inflammation in humans [69]. Previous transcriptomic studies in marine invertebrates have shown differential regulation of *Dmbt1* expression in response to both bacterial challenge and symbiont acquisition [70–73], supporting the hypothesis that this gene contributes to maintaining healthy associations with commensal microbes [73]. Downregulation of this gene in SCTLD-affected corals was accompanied by shifts in mucus microbial community structure [33,48]. The higher expression of *Dmbt1* in apparently healthy coral colonies suggests a role in disease resistance through the establishment and maintenance of mucosal microbial homeostasis and the prevention of opportunistic infection.

The top feature in HH *C. goreaui* annotated to Protein NO VEIN (*NOV*), a plant-specific nuclear factor required for cell-fate decisions in *Arabisopsis thaliana* [74]. Interpro protein domain analysis found a Histidine kinase/Heat Shock Protein 90 (HSP90)-like ATPase superfamily domain within this gene’s protein-coding sequence. This conserved domain is found in proteins involved in both cellular signaling and protein folding through the histidine kinase domain and the HSP90-like ATPase domain, respectively [61]. Histidine kinases are responsible for triggering signal transduction in response to environmental stimulus, such as nutrient availability, pH, osmolarity and light, while HSP90-like ATPases are molecular chaperones that play an essential role in protein folding and stabilization [61]. High expression of this gene in *C. goreaui* from apparently healthy coral tissue may therefore suggest heightened sensitivity to environmental cues and prevent the accumulation of misfolded proteins. Additional coral disease research may benefit from elucidating the precise function of this gene in *C. goreaui*, given its putative ability to provide natural resilience against SCTLD and its role in cellular homeostasis.

### HD Tissue Is Responding to SCTLD by Promoting Symbiont Acquisition and Retention

Visibly healthy tissue on SCTLD-affected colonies has been shown to exhibit cellular morphological changes associated with Symbiodiniaceae pathology, such as chloroplast deformity and cavity formation [39]. While the apparently healthy tissue on a diseased colony can have intermediate levels of expression of many genes (Fig 4 and 6), our supervised ML approach identified features strongly up- or downregulated by both *M. cavernosa* and *C. goreaui* in this tissue. Our results provide further evidence that apparently healthy tissue on a diseased colony exhibits a unique gene expression profile indicative of a preemptive response to infection.

In contrast to the high levels of fatty acid metabolism seen in the HH tissue enrichments, functional enrichment of the HD features showed significant upregulation of amide biosynthesis in both *M. cavernosa* and *C. goreaui*. These results are indicative of a shift in metabolic functioning between symbiotic partners during the onset of colony infection with SCTLD. Amides serve as components of proteins, enzymes, and signaling molecules and are involved in many biological processes. In response to stress or infection, the cell may increase amide biosynthesis to ensure an adequate supply of amino acids for the synthesis of stress-response proteins or for repairing damaged proteins [75]. This upregulation of amide biosynthesis by both the coral host and its algal endosymbiont provides evidence that the apparently healthy tissue on a SCTLD-affected colony exhibits unique metabolic signatures, perhaps in preparation to mount an immune response.

Analysis of the top features from HD *M. cavernosa* provides further evidence of a unique response to SCTLD infection in the apparently healthy tissue of a diseased coral colony. Within those features, we found upregulation of *Rab5a*, a gene whose protein product is a prototypical marker of early endosomal and phagocytic vesicles [76]. In cnidarians, Rab proteins play a crucial role in mediating endosymbiosis through the maturation or arrest of Symbiodiniaceae-hosting phagocytic vesicles called symbiosomes [77]. In the sea anemone *Aiptasia pulchalla* (*Exaiptasia diaphana*), Rab5 localizes to symbiosomes containing newly ingested or established Symbiodiniaceae, but not in symbiosomes containing dead or dysfunctional Symbiodiniaceae [78], highlighting its role in arresting phagosome maturation and subsequent endosymbiont digestion. Interestingly, some intracellular pathogens have been shown to modulate host *Rab5* expression, allowing them to persist and replicate within arrested phagosomes [79–81]. The upregulation of *Rab5a* by *M. cavernosa* in the HD tissue may therefore indicate an effort, either mediated by the coral host or its Symbiodiniaceae, to promote and maintain algal endosymbiont retention within the apparently healthy tissue of a colony affected with SCTLD.

In addition, HD *M. cavernosa* exhibited a downregulation of two genes (*cfap298* and *fam166b*) involved in cilia movement and assembly. In corals, cilia are directly involved in heterotrophic feeding and nutrient acquisition [82], and the downregulation of *cfap298* and *fam166b* may indicate a reduction in host nutrient acquisition from the surrounding seawater. Interestingly, within the top HD features from *C. goreaui*, we saw upregulation of *STY17*, a serine/threonine protein kinase that phosphorylates chloroplast-destined precursor proteins. In the plant *Arabidopsis thaliana*, *STY17* knockout resulted in delayed assimilation of chlorophyll and a reduction in photosynthetic capacity, highlighting the vital role of this gene in chloroplast biogenesis and differentiation [83]. A shift in the metabolic exchange between corals and their algal endosymbionts in the apparently healthy tissue on a SCTLD-affected colony likely indicates an increased reliance on Symbiodiniaceae for energy and nutrient acquisition.

### LD Tissue is characterized by immune activity, loss of extracellular matrix structure, and host-endosymbiont dysbiosis

Functional enrichment of features in LD *M. cavernosa* shows upregulation of genes involved in innate immunity, notably C-type lectin (CTL) receptor binding and NF-κB signaling pathways. CTLs, identified in many cnidarian species [84], recognize exogenous ligands to activate innate immunity via NF-κB and lectin complement pathways [85]. When activated, NF-κB launches a wide array of host responses, including phagocytosis, inflammation, and antimicrobial mechanisms, often promoting cell survival and inhibiting apoptosis [86]. Previous studies have shown that Symbiodiniaceae are able to suppress host NF-κB signaling to maintain symbiosis within their cnidarian hosts [87–90]. However, during pathogenic stress, this suppression may be disrupted. For example, symbiotic *Cassiopea xamachana* exposed to *Serratia marcescens* exhibited reduced survival rates compared to their aposymbiotic counterparts, alongside NF-κB upregulation that was not seen in the aposymbiotic group [91]. In addition, corals exposed to SCTLD *ex situ* exhibited increased expression of genes involved in NF-κB signaling, but this response was absent following amoxicillin treatment that effectively slowed or halted lesion progression [50]. These findings suggest that direct pathogen exposure triggers NF-κB activation in symbiotic cnidarians, leading to inflammation that can exacerbate tissue damage if left unregulated. It therefore appears that the ability to appropriately regulate NF-κB signaling and maintain symbiotic homeostasis is crucial in arresting lesion progression.

Additionally, we saw enrichment of the term ‘autophagosome maturation’ within the LD *M. cavernosa* features, suggesting a heightened activation of autophagy at the lesion tissue in corals affected with SCTLD. One of the most ancient forms of innate immunity in animal cells, autophagy, is the process by which cytoplasmic contents, including intracellular bacteria and viruses, are engulfed within a double-membrane autophagosome which is then delivered to lysosomes for degradation and recycling [92]. Autophagy of Symbiodiniaceae is called symbiophagy, the process by which the host-derived symbiosome is transformed from an arrested state of phagocytosis into a digestive organelle [93]. Rab7, an established marker of symbiophagy, was upregulated in multiple species in response to SCTLD infection, implicating symbiophagy in the pathology of SCTLD [48]. While we did not find *Rab7* in our top LD *M. cavernosa* features, the enrichment of autophagosome maturation is further evidence of either an activation of an autophagic immune response against a pathogen, an activation of symbiophagy in response to dead or dysfunctional Symbiodiniaceae, or both concomitantly. The significance of autophagosome maturation in SCTLD-affected corals further highlights the intricate relationship between innate immunity and symbiosis maintenance during the pathogenesis of SCTLD and represents a promising target for further research in disease prevention and intervention efforts.

Analysis of the top LD features in *M. cavernosa* shows that SCTLD lesion tissue is predominantly characterized by a dramatic downregulation of genes involved in maintaining extracellular matrix (ECM) and actin cytoskeletal structure. The top feature in LD *M. cavernosa* was the downregulated *Tpm1*, a gene implicated in stabilizing cytoskeletal actin filaments in non-muscle cells. Interestingly, Beavers et al. 2023 showed that a similar gene, Tropomyosin 4 (*Tpm4*), exhibited significant downregulation in five species of coral after experimental exposure to SCTLD [48]. Furthermore, tropomyosin is also downregulated in a closely related scleractinian coral, *Orbicella faveolata,* during thermal stress and bleaching [94]. It is therefore likely that actin cytoskeletal structure in SCTLD lesion tissue is experiencing disruption due to oxidative stress and dysregulated nutrient transfer caused by symbiosis breakdown.

In addition, five collagen genes (two Collagen alpha-1(II) chain homologs (*col2a1 [a]* and *col2a1 [b],* two Collagen alpha-2(I) chain homologs (*col1a2 [a]* and *col1a2 [b]*, and one collagen alpha-2(XI) chain (*col11a1*)) were downregulated within the top LD *M. cavernosa* features. These results contrast those seen in a SCTLD transmission experiment by Traylor-Knowles et al. 2021, who saw differential up- and down-regulation of collagen genes in response to early infection, leading to their hypothesis that corals affected with SCTLD are activating wound healing mechanisms in response tissue degradation [49]. In our field-collected samples, however, the drastic downregulation of ECM components indicates a compromised ability of the coral tissue to maintain structural integrity, thereby hindering effective lesion healing and potentially increasing vulnerability to secondary infections. These differences in ECM gene expression dynamics could stem from temporal variations in sampling conditions: Traylor-Knowles et al., (2021) likely captured early-stage responses during disease onset, whereas our samples were collected from a natural reef environment, where the date of SCTLD onset was unknown. Understanding these variations in ECM and cytoskeletal dynamics throughout different temporal stages of SCTLD progression may therefore be crucial for the development of targeted treatment strategies and conservation efforts aimed at combating this deadly disease.

Examining host and endosymbiont features together, we find evidence of photosystem dysfunction, ROS production, and dysbiosis in the SCTLD lesion tissue. *EDL1*, a glycosyltransferase-like protein that acts as a negative regulator of photomorphogenesis, was upregulated in *C. goreaui* within the lesion tissue, signifying a redirection of cellular resources away from growth-related processes. In addition, *FCPE*, a component of the light-harvesting complex embedded in the thylakoid membrane, was also upregulated in *C. goreaui* from the lesion tissue, which could represent a strategy to enhance the efficiency of photosynthesis under stress conditions. However, within the top features in LD *M. cavernosa*, we found downregulation of *GFPL*, a green fluorescent protein-like pigment that is implicated in photoprotection of Symbiodiniaceae [95] as well as a drastic increase in expression of *sodA,* an established coral antioxidant involved in reducing harmful superoxide radicals (Barshis et al. 2013; Lesser 1997). A compromise in *M. cavernosa’s* ability to protect its algal endosymbiont from excess light in the lesion tissue could explain the observed shifts in chloroplast function by *C. goreaui*, leading to photosystem stress, leakage of ROS into host tissue, and a subsequent antioxidant response by the coral host. These gene expression signatures resemble those seen during bleaching [98] and represents further evidence of host-endosymbiont dysbiosis within the SCTLD lesion tissue.

### Novel application of feature selection for characterizing coral health states

Coral research has inherent challenges, the most limiting being the low availability of samples from vulnerable or threatened populations and furthermore, the low availability of these samples exhibiting active pathological signs of a specific disease or health state. While it is preferable to use large sample sizes to draw robust conclusions in gene expression studies, it is often not possible. Keeping this in mind, the methodology presented in this study leverages a supervised ML-based feature selection algorithm and validation procedure that is designed to identify important gene expression signatures even from limited sample sizes.

We selected sigFeature as our feature selection algorithm because of its ability to address the challenges posed by small-sample gene expression studies [52]. By combining SVM-RFE–which excels at handling high-dimensional data [99]–with the differential expression *t*-statistic to rank genes, sigFeature ensures that selected features are both discriminative and biologically meaningful. This approach allows us to uncover important gene expression patterns that traditional methods might miss. This methodology aligns with emerging approaches that aim to combine ML techniques with biological knowledge to enhance biomarker discovery [100]. Additionally, the ability of sigFeature to handle unbalanced class distributions and high-dimensional data makes it well-suited for coral gene expression studies. Furthermore, the feature selection process was rigorously evaluated using external stratified 10-fold cross-validation, a standard technique for assessing classifier model performance [101,102]. This cross-validation method operates independently of the feature selection process and provides unbiased performance metrics across all folds, offering a reliable estimate of feature classification importance. In this study, cross-validation results were consistent with those reported in larger dataset studies [52] and exhibited stable performance across all tissue health states.

## Conclusion

Our gene expression profiling of various tissue health states in *M cavernosa* and its dominant algal endosymbiont, *C. goreaui*, supports evidence that SCTLD causes dysbiosis between the coral host and its Symbiodiniaceae. While this study does not confirm the etiologic agent of SCTLD, it does highlight the unique shifts in host-endosymbiont functioning both during the onset of colony infection and during lesion progression. Apparently healthy tissue on SCTLD-affected colonies appears to be mounting a response to colony infection by promoting Symbiodinaiceae uptake and retention, possibly to increase autotrophic nutrient acquisition to prepare an immune response. If SCTLD is caused by a pathogen of Symbiodinaiceae, this strategy would be detrimental to the coral host as it could inadvertently increase the acquisition of affected endosymbionts. Lesion tissue, alternatively, showed strong signals of runaway inflammation, loss of cellular integrity, and an inability to maintain symbiotic homeostasis. Some of the gene expression signatures in the lesion tissue resemble bleaching responses, such as disruption of actin cytoskeleton structure and oxidative stress in the coral and photosystem dysregulation in the algal endosymbiont. The compounded stress of mounting an inflammatory response in addition to a breakdown in host-endosymbiont physiology could therefore explain the rapid tissue loss observed in SCTLD progression.

While our research offers important insights into SCTLD disease mechanisms, there are some limitations that should be considered. Focusing on a single coral species and Symbiodiniaceae association from the U.S. Virgin Islands means our results may not capture the dynamics present in other coral species and regions affected by SCTLD. Additionally, our sample size was limited due to the availability of suitable coral colonies. Despite this, the use of SVM-RFE allowed us to drive meaningful insights from the available data, and consistency across samples within the same tissue healthy state support our findings’ validity. Future studies should aim to incorporate multiple coral species and larger sample sizes to provide a more comprehensive understanding of SCTLD pathogenesis and its impact on coral-symbiont functioning.

Our bioinformatic pipeline utilizing a supervised ML feature selection algorithm offers a novel method to characterize disease pathogenesis in corals and their algal endosymbionts by isolating the most biologically relevant genes with high classification power. With the increasing availability of high-throughput data, these methods can be integrated into omics analyses to accurately characterize various coral physiological states, such as white plague disease, bleaching, and emerging disease outbreaks. With this feature selection framework, we can better understand the health states of endangered coral species to develop effective and long-lasting management efforts.

## METHODS

### Sample collection and design

All samples were collected under permit #DFW19057U authorized by the Department of Planning and Natural Resources Coastal Zone Management. Coral fragments approximately 5cm in diameter were collected by divers on SCUBA with hammers and chisels from two reefs in St. Thomas, United States Virgin Islands (USVI) showing signs of active SCTLD in February of 2020: Buck Island (18.27883°, −64.89833°) and Black Point (18.3445°, −64.98595°). Black Point, a nearshore reef, first exhibited SCTLD cases between December 2018 and January 2019, as documented in. Buck Island, situated near an offshore undeveloped island, recorded its first cases of SCTLD in October 2019, also documented in. Current environments at both sites are similar. SCTLD-affected corals were identified based on displaying acute multifocal lesions consistent with the SCTLD case definition [103]. Lesions were bright white where the skeleton had recently been denuded of tissue, with no visible algal colonization at the skeletal/tissue boundary, indicating actively expanding lesions.

At both sites, one coral fragment was collected from each apparently healthy colony (Buck Island, *n=3*; Black Point, *n*=3), termed apparently healthy tissue on an apparently healthy colony (HH). Two fragments were collected from each diseased colony: one immediately adjacent to the SCTLD lesion line (Buck Island, *n=*3; Black Point, *n=*5), termed lesion tissue on a diseased colony (LD), and one as far away from the lesion line as possible (approximately 10 cm from the lesion line) (Buck Island, *n=*3; Black Point, *n=*5), termed apparently healthy tissue on a diseased colony (HD). The sampling scheme (shown in Fig 1A-C) aimed to capture the variability in gene expression across different tissue health states while ensuring consistency in sampling methodology. Coral fragments were placed in individual bags that were sealed and transported to land on ice before being flash frozen at −80°C.

### RNA extraction and sequencing

Total RNA was extracted from all 22 coral fragments following the protocol outlined in [48] using the RNAqueous-4PCR Total RNA Isolation Kit from Invitrogen (Life Technologies AM1914). About one gram of frozen coral tissue was scraped off each fragment into a 2 mL microcentrifuge tube using a sterilized bone cutter. Lysis buffer was added to each microcentrifuge tube followed by mechanical disruption using a refrigerated Qiagen Tissuelyser II at 30 oscillations/s for 60 s. Elution was performed in two 30 µL steps at a time. After combining elutions, contaminating DNA and chromatin were removed using the Ambion DNase I kit from Invitrogen (Life Technologies AM 2222). Resulting total RNA samples were sent to Novogene Co., LTD (Beijing, China) for quality assessment using an Agilent Bioanalyzer 2100. All samples passed quality assessment with RNA integrity (RIN) values ≥ 7 and were preprocessed for mRNA enrichment using polyA tail capture. cDNA libraries were prepared using the NEBNext Ultra II RNA Library Prep Kit from Illumina and sequenced on the Illumina NovaSeq 6000 for 150 bp, paired-end sequencing.

### M. cavernosa transcriptome assembly and annotation

All bioinformatic analyses were carried out on the Frontera system of the Texas Advanced Computing Center [104]. Raw reads from Novogene were adapter-trimmed and quality-filtered in one step using FastP v. 0.20.1 [105] using the -c flag for base correction and the -x flag for polyA tail trimming. Then, six samples (two from each health state selected at random) were used to generate a *de novo* metatranscriptome using Trinity v. 2.14.0 [106]. Non-coral transcripts were filtered out of this metatranscriptome using the *in-silico* filtration method outlined previously [107]. First, the longest transcript isoform was obtained using the get_longest_isoform_seq_pr_trinity_gene.pl script within the Trinity v. 2.14.0 package [106]. This assembly was then Blasted against a Master Coral database [108] comprised of both genome-derived predicted gene models and transcriptomes spanning a wide diversity of coral families using BlastX v. 2.2.27 [109]. Transcripts with less than 95% identity of this Master Coral database and shorter than 150 bp in length were filtered out of the assembly. TransDecoder v. 5.5.0 [110] was used to generate a predicted protein-coding sequence from the longest open reading frame from each transcript, resulting in a predicted proteome for *M. cavernosa*. Sequences with high similarity were then collapsed using cd-hit v. 4.8.1 [111] under default parameters. The resulting sequences were then extracted from the initial assembly to generate the *M. cavernosa* reference transcriptome. Assembly metrics can be found in Table 1. Finally, the transcriptome was annotated with reviewed UniprotKB/Swiss-Prot Entry IDs using BlastX v. 2.2.27 [109].

### Isolation and quantification of holobiont reads

To separate *M. cavernosa* and Symbiodiniaceae reads, BBSplit v. 38.90 [60] was used with the *M. cavernosa* reference transcriptome generated above and Symbiodiniaceae transcriptomes of similar assembly quality representing the genera *Symbiodinium, Breviolum, Cladocopium,* and *Durusdinium* sourced from previous publications as binning references [58,112–114]. The binning statistics output was used to identify *Cladocopium* as the dominant Symbiodiniaceae genera present in each sample (Table S1). A predicted proteome was generated from the *C. goreaui* reference transcriptome used above with TransDecoder v. 5.5.0 [110]. Similar sequences in the proteome were collapsed using cd-hit v. 4.8.1 [111]. The resulting sequences were then extracted from their initial assembly to generate the final *C. goreaui* reference transcriptome. Assembly metrics for this transcriptome can be found in Table 1. The resulting *C. goreaui* assembly was then annotated with reviewed UniprotKB/Swiss-Prot Entry IDs using BlastX v. 2.2.27 [109]. Finally, *M. cavernosa* and *C. goreaui* reads from all 22 samples were mapped to their respective transcriptome and quantified using Salmon v. 1.5.2 [115] under default parameters for *M. cavernosa* reads and a kmer value of 23 for *C. goreaui* reads.

### Feature selection on holobiont gene expression

All data analysis was performed using R v. 4.2.2 [116]. Gene count matrices were generated for the quantified *M. cavernosa* and *C. goreaui* transcripts using Tximport v. 1.16.1 [117]. Annotated *M. cavernosa* and *C. goreaui* genes with an evalue < 1.0e^−6^ were kept for differential expression analyses. Regularized log (rlog) normalized expression was obtained in each dataset using DESeq2 v. 1.38.3 [118] with the design *∼Site + Disease_state,* removing genes with an average rlog expression < 10. Prior to feature selection, a Principal Component Analysis (PCA) was performed to assess gene expression differences between the endemic and epidemic sites, and no obvious separation between sites was identified (Fig S1). Therefore, all samples belonging to the same health state (HH, HD, or LD) were pooled across sites, resulting in the following sample sizes per health state: HH: *n* = 6, HD: *n =* 8, and LD: *n =* 8. Then, the *sigFeature()* feature selection algorithm within the sigFeature v. 1.16.0 R package [52] was used to produce a ranked list of genes for each health state in *M. cavernosa* and *C. goreaui* (Fig 1D) (Tables S2-7). This algorithm combines support vector machine recursive feature elimination (SVM-RFE) with statistical significance testing (*t-*statistic), ranking genes by their classification accuracy and their differential expression levels to ensure biological relevance [100].

To evaluate the classification efficacy of features from each tissue health state, we performed external stratified 10-fold cross validation on the top 400 features from each dataset using the *sigFeature.enfold()* [52] function. This function partitions the dataset (22 samples) into 10 approximately equal-sized folds (two folds with three samples each, and eight folds with two samples each) and ranks the features in each fold based on their contribution to class separation through recursive elimination. For each iteration, nine folds were used for feature selection and training, while the remaining fold was reserved for testing the performance of the selected features. This procedure was repeated 10 times, with each fold serving as the validation set exactly once, ensuring that every sample in the dataset was used for both training and validation.

Following the cross-validation, we applied the *sigFeatureFrequency()* [52] function to rank features based on their frequency of selection across all ten iterations. This function generated a ranked list of the top 400 features, assigning frequency scores based on how often each feature appeared in the top 400 features of each iteration of cross-validation. We then used these frequency scores to perform a feature sweep analysis using the *sigCVError()* [52] function, which calculates the cross-validation error rates across increasing numbers of the top features (from 1 to 400). The error rates were plotted to visualize both the mean error and the standard deviation across the 10 iterations of cross-validation (Fig 1E-F).

### Functional enrichment of top features in M. cavernosa

Functional enrichment was performed on both the up- and down-regulated genes within the top 500 features from each tissue health state in *M. cavernosa* and *C. goreaui*. To find the upregulated features in each tissue health state, log_2_FoldChange values from DESeq2 relative to the two other tissue health states were obtained for the top 500 features. Features with a log_2_FoldChange > 0 relative to the two other tissue health states were classified as “upregulated”, while those with a log_2_FoldChange < 0 were classified as “downregulated” (Tables S8-13). For example, to identify upregulated HH features, log_2_FoldChange was obtained for the comparisons between HH and HD corals as well as between HH and LD corals, and those features out of the top 500 HH features that had higher expression in HH corals relative to both HD and LD corals were deemed “upregulated”. To perform functional enrichment of those up- and down-regulated features, first the predicted proteomes of *M. cavernosa* and *C. goreaui* generated above were uploaded to STRING v. 12.0 [119] using the ‘Add organism’ tool (*M. cavernosa* STRING ID: STRG0A38MOJ, *C. goreaui* STRING ID: STRG0A06ZQW). Then, lists of the up-regulated and down-regulated features were uploaded to STRING separately. All Gene Ontology, Reactome, and KEGG terms were saved, and a curated list of the 5 most relevant and non-redundant up- and downregulated terms were plotted (Fig 2 and 3).

### Expression analysis of potential tissue health state biomarkers

The top 15 features from each tissue health state in both *M. cavernosa* and *C. goreaui* were selected for further analysis due to their high classification potential. Rlog values were obtained for each list of the top 15 features in *M. cavernosa* and plotted in relative expression heatmaps. To confirm gene function, protein domain architecture was obtained and analyzed for each feature by uploading their protein sequence to InterPro [61] (Tables S14-19). From each tissue health state’s top 15 features, the top feature as well as four other highly relevant features were plotted in boxplots (Fig 4B, 5B, and 6B). Additionally, three features from each tissue health state in *C. goreaui* were selected based on their differential expression significance and functional relevance and were plotted in boxplots (Fig 7).

## ACKNOWLEDGEMENTS

The authors would like to thank the facilities and staff at the University of the Virgin Islands Center for Marine and Environmental Studies (CMES), and Dr. Cynthia Becker, for their assistance with field collections. Samples were collected under permit #DFW19057U authorized by the Department of Planning and Natural Resources Coastal Zone Management. This is CMES contribution #256. Bioinformatic analyses were performed on the Frontera supercomputer at the Texas Advanced Computing Center. This work was funded by National Science Foundation (NSF) VI EPSCoR 0814417 and 1946412 and NSF (Biological Oceanography) award numbers 1928753 to M.E.B., 1928771 to L.D.M., as well as 1928761 and 1938112 to A.A., NSF EEID award number 2109622 to M.E.B., A.A., L.D.M., and a NOAA OAR Cooperative Institutes award to A.A. (#NA19OAR4320074).

---

**Fig S1.**
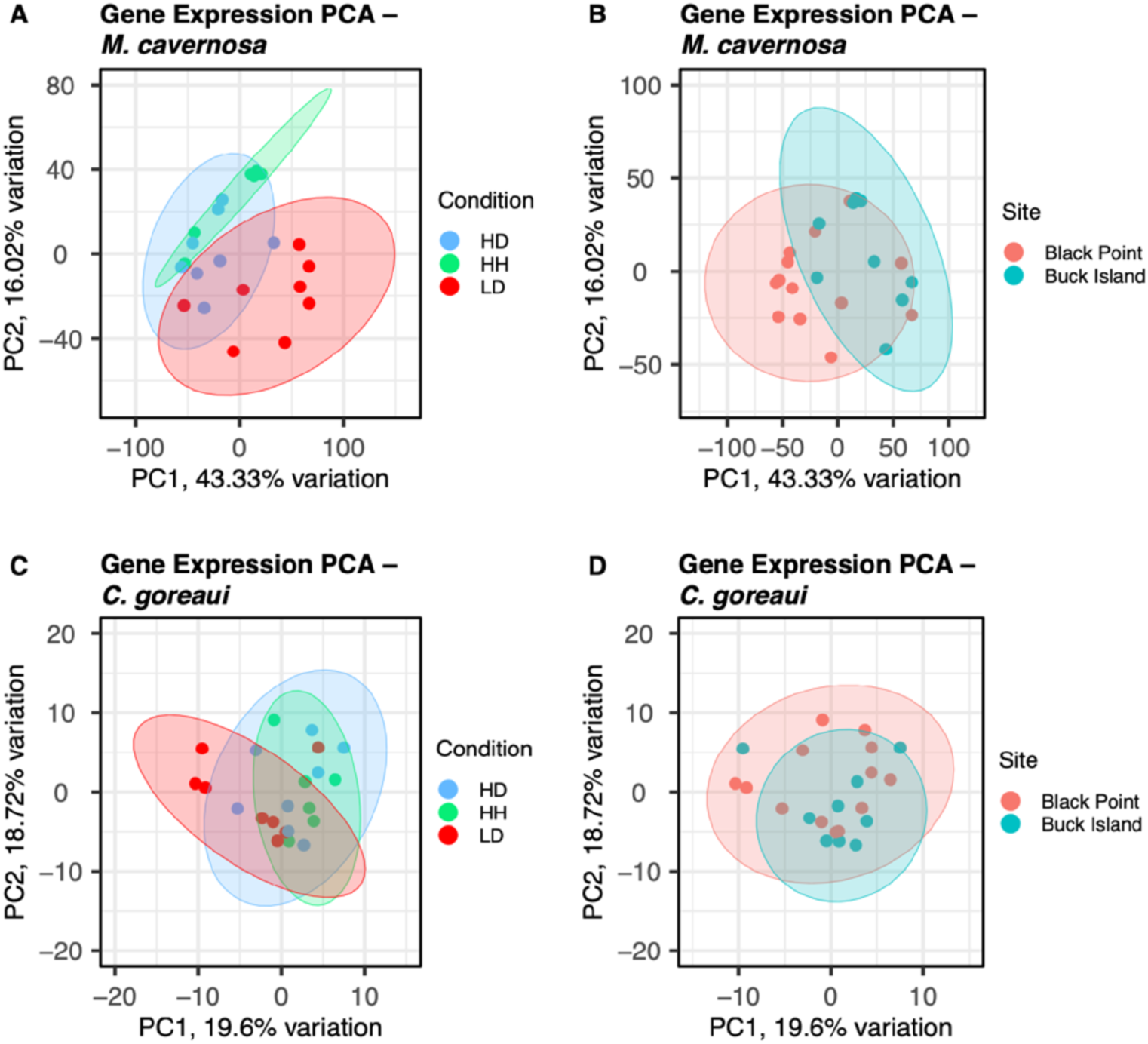
Principal Component Analysis (PCA). PCAs showing the regularized log (rlog) expression of all *M. cavernosa* genes grouped by (A) tissue health state and (B) collection site, and all *C. goreaui* genes grouped by (C) tissue health state and (D) collection site.

**Fig S2.**
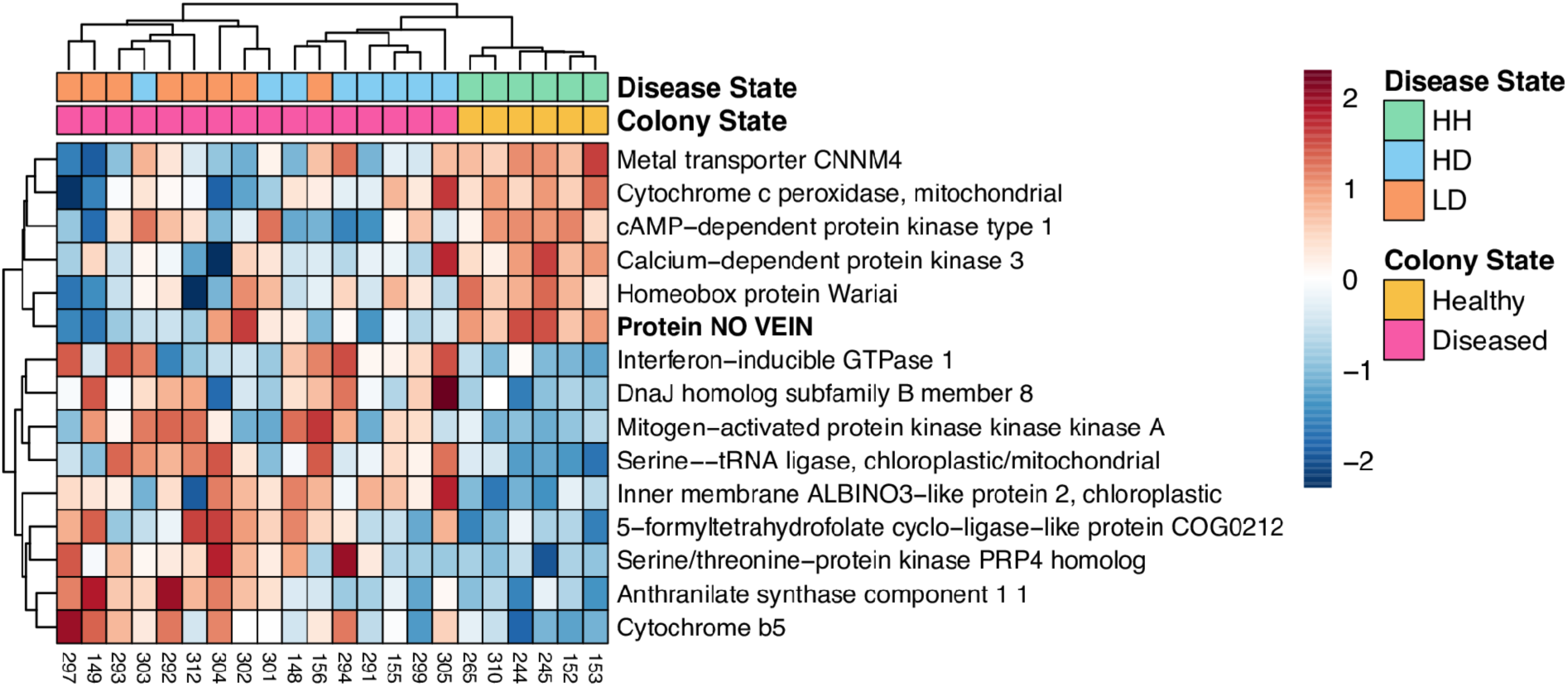
Relative expression heatmap of the top 15 HH features from *C. goreaui*. Red boxes signify elevated expression relative to the row mean, and blue boxes signify lowered expression relative to the row mean. The top feature is shown in bold.

**Fig S3.**
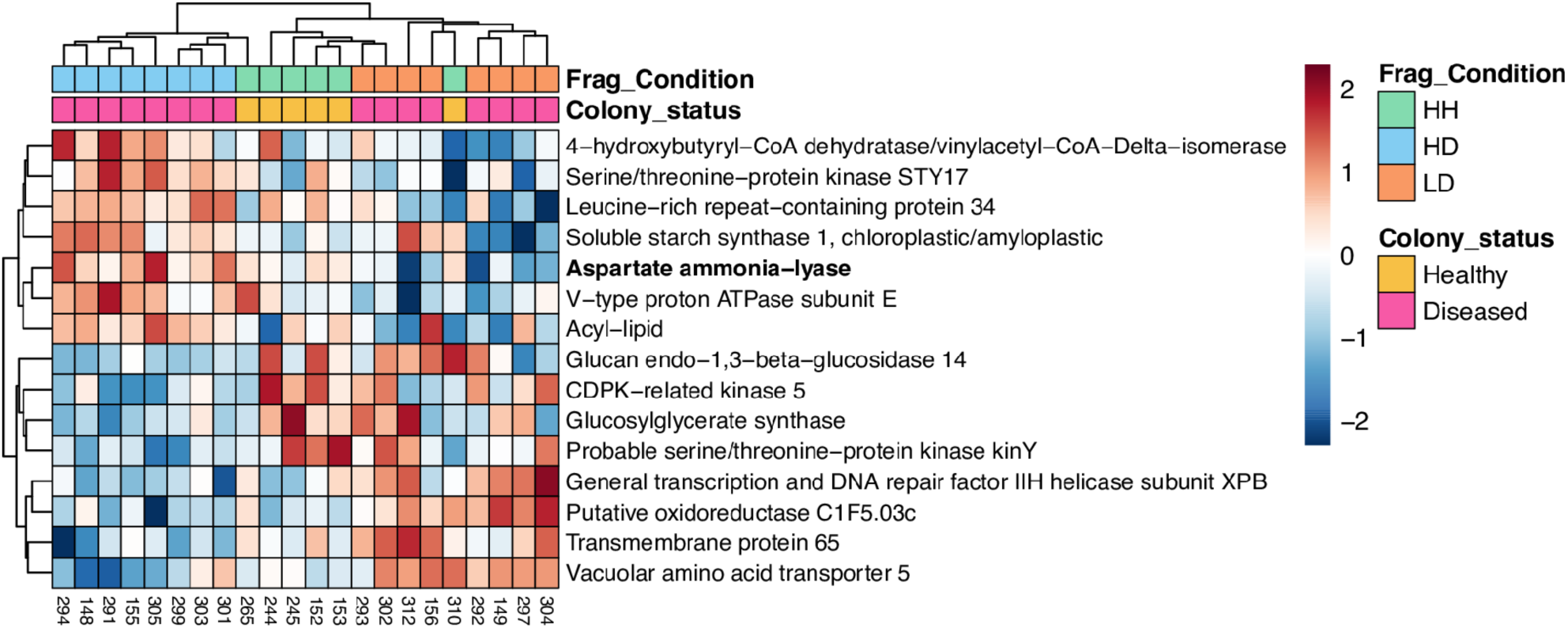
Relative expression heatmap of the top 15 HD features from *C. goreaui*. Red boxes signify elevated expression relative to the row mean, and blue boxes signify lowered expression relative to the row mean. The top feature is shown in bold.

**Fig S4.**
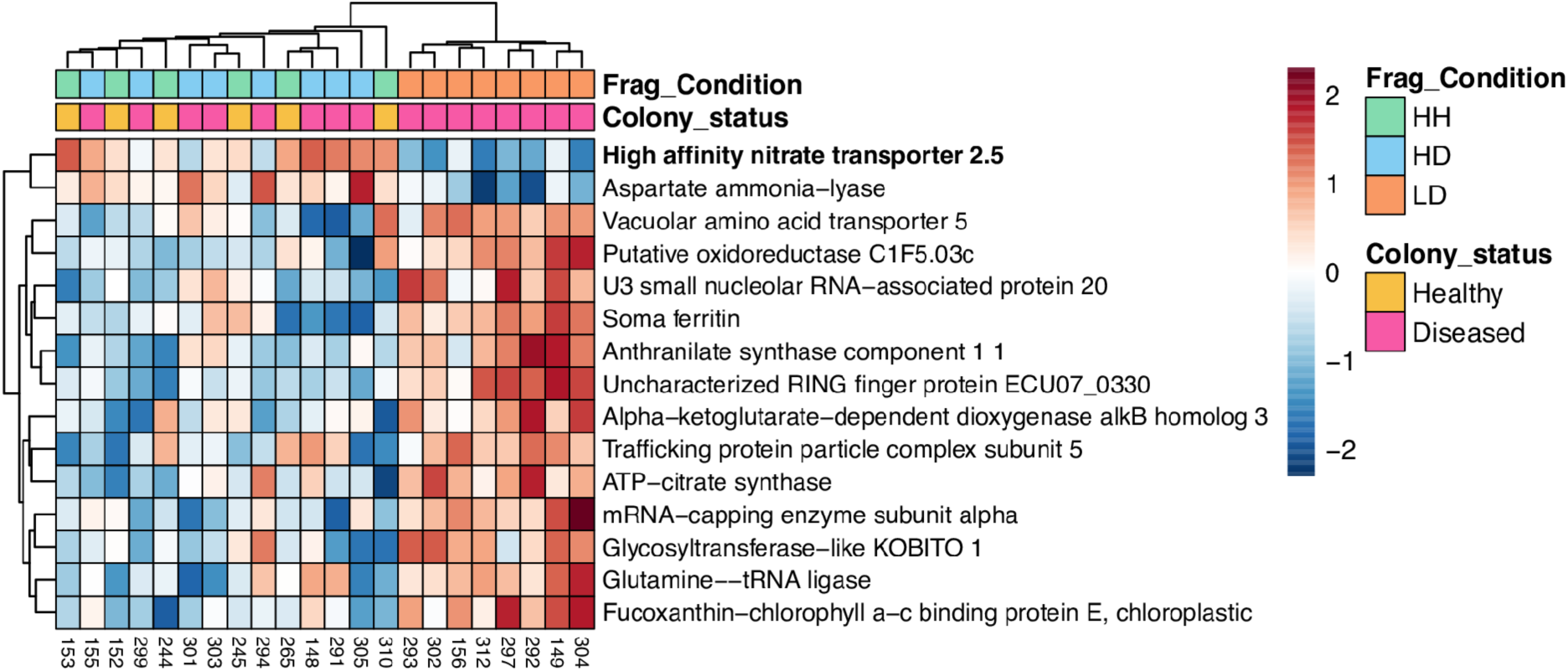
Relative expression heatmap of the top 15 LD features from *C. goreaui*. Red boxes signify elevated expression relative to the row mean, and blue boxes signify lowered expression relative to the row mean. The top feature is shown in bold.

## DATA AVAILABILITY STATEMENT

All raw sequence data and associated sample metadata are deposited in the NCBI SRA database (BioProject PRJNA1062758). Publicly available data used in this study include the reference transcriptomes for *Symbiodinium* CassKB8 (BioProject PRJNA80085), *Breviolum minutum* (BioProject PRJNA274852), *Cladocopium goreaui* (BioProject PRJNA307543), and *Durusdinium trenchii* (BioProject PRJNA508937). The Master Coral database used is available in a public Zenodo repository (https://doi.org/10.5281/zenodo.7838980) (Beavers 2023). Additional sampling information and all code used in this study are available on GitHub at https://github.com/kbeavz/SCTLD-Feature-Selection-USVI/tree/1.1.0 and we used Zenodo to assign a DOI to the repository (DOI: <u>10.5281/zenodo.12744785</u>).

## AUTHOR CONTRIBUTIONS

Conceptualization: K.M.B., E.V.B., A.A., L.D.M. Sample collection: K.M.B., M.E.B., A.A. RNA extraction and processing: K.M.B. Data analysis: K.M.B, D.G.A., with input from M.E. and E.V.B. Manuscript writing: K.M.B and L.D.M., with editing contributions from all authors.

## REFERENCES

1. Vega Thurber R, Mydlarz LD, Brandt M, Harvell D, Weil E, Raymundo L, et al. Deciphering Coral Disease Dynamics: Integrating Host, Microbiome, and the Changing Environment. Front Ecol Evol. 2020;8. doi:10.3389/fevo.2020.575927

2. Sully S, Hodgson G, van Woesik R. Present and future bright and dark spots for coral reefs through climate change. Glob Change Biol. 2022;28: 4509–4522. doi:10.1111/gcb.16083

3. Eddy TD, Lam VWY, Reygondeau G, Cisneros-Montemayor AM, Greer K, Palomares MLD, et al. Global decline in capacity of coral reefs to provide ecosystem services. One Earth. 2021;4: 1278–1285. doi:10.1016/j.oneear.2021.08.016

4. Bruno JF, Selig ER. Regional Decline of Coral Cover in the Indo-Pacific: Timing, Extent, and Subregional Comparisons. PLOS ONE. 2007;2: e711. doi:10.1371/journal.pone.0000711

5. Hughes TP, Anderson KD, Connolly SR, Heron SF, Kerry JT, Lough JM, et al. Spatial and temporal patterns of mass bleaching of corals in the Anthropocene. Science. 2018;359: 80–83. doi:10.1126/science.aan8048

6. Hoegh-Guldberg O, Bruno JF. The Impact of Climate Change on the World’s Marine Ecosystems. Science. 2010;328: 1523–1528. doi:10.1126/science.1189930

7. Hughes TP, Kerry JT, Baird AH, Connolly SR, Dietzel A, Eakin CM, et al. Global warming transforms coral reef assemblages. Nature. 2018;556: 492–496. doi:10.1038/s41586-018-0041-2

8. Boström-Einarsson L, Babcock RC, Bayraktarov E, Ceccarelli D, Cook N, Ferse SCA, et al. Coral restoration – A systematic review of current methods, successes, failures and future directions. PLOS ONE. 2020;15: e0226631. doi:10.1371/journal.pone.0226631

9. Omori M. Coral restoration research and technical developments: what we have learned so far. Mar Biol Res. 2019;15: 377–409. doi:10.1080/17451000.2019.1662050

10. Quigley KM, Baird AH. Future climate warming threatens coral reef function on World Heritage reefs. Glob Change Biol. 2024;30: e17407. doi:10.1111/gcb.17407

11. Vollmer SV, Selwyn JD, Despard BA, Roesel CL. Genomic signatures of disease resistance in endangered staghorn corals. Science. 2023;381: 1451–1454. doi:10.1126/science.adi3601

12. Quigley KM, van Oppen MJH. Predictive models for the selection of thermally tolerant corals based on offspring survival. Nat Commun. 2022;13: 1543. doi:10.1038/s41467-022-28956-8

13. Mcleod E, Anthony KRN, Mumby PJ, Maynard J, Beeden R, Graham NAJ, et al. The future of resilience-based management in coral reef ecosystems. J Environ Manage. 2019;233: 291–301. doi:10.1016/j.jenvman.2018.11.034

14. van Oppen MJH, Oliver JK, Putnam HM, Gates RD. Building coral reef resilience through assisted evolution. PNAS. 2015;112: 2307–2313. 10.1073/pnas.1422301112

15. Kleypas J, Allemand D, Anthony K, Baker AC, Beck MW, Hale LZ, et al. Designing a blueprint for coral reef survival. Biol Conserv. 2021;257: 109107. doi:10.1016/j.biocon.2021.109107

16. Quigley KM, Donelson JM. Selective Breeding and Promotion of Naturally Heat-Tolerant Coral Reef Species. 2nd ed. Oceanographic Processes of Coral Reefs. 2nd ed. CRC Press; 2024.

17. Alvarez-Filip L, González-Barrios FJ, Pérez-Cervantes E, Molina-Hernández A, Estrada-Saldívar N. Stony coral tissue loss disease decimated Caribbean coral populations and reshaped reef functionality. Commun Biol. 2022;5: 1–10. doi:10.1038/s42003-022-03398-6

18. Estrada-Saldívar N, Quiroga-García BA, Pérez-Cervantes E, Rivera-Garibay OO, Alvarez-Filip L. Effects of the Stony Coral Tissue Loss Disease Outbreak on Coral Communities and the Benthic Composition of Cozumel Reefs. Front Mar Sci. 2021;8. doi:10.3389/fmars.2021.632777

19. Hayes NK, Walton CJ, Gilliam DS. Tissue loss disease outbreak significantly alters the Southeast Florida stony coral assemblage. Front Mar Sci. 2022;9. Available: https://www.frontiersin.org/articles/10.3389/fmars.2022.975894

20. Heres MM, Farmer BH, Elmer F, Hertler H. Ecological consequences of Stony Coral Tissue Loss Disease in the Turks and Caicos Islands. Coral Reefs. 2021;40:609–624. doi:10.1007/s00338-021-02071-4

21. Walton CJ, Hayes NK, Gilliam DS. Impacts of a Regional, Multi-Year, Multi-Species Coral Disease Outbreak in Southeast Florida. Front Mar Sci. 2018;5. doi:10.3389/fmars.2018.00323

22. Kramer PR, Roth L, Lang J. Map of stony coral tissue loss disease outbreak in the Caribbean. 2019. Available: www.agrra.org

23. NOAA. Stony Coral Tissue Loss Disease Case Definition. 2018. Available: https://floridadep.gov/sites/default/files/Copy%20of%20StonyCoralTissueLossDisease_CaseDefinition%20final%2010022018.pdf

24. Precht WF, Gintert BE, Robbart ML, Fura R, van Woesik R. Unprecedented Disease-Related Coral Mortality in Southeastern Florida. Sci Rep. 2016;6: 31374. doi:10.1038/srep31374

25. Alvarez-Filip L, Estrada-Saldívar N, Pérez-Cervantes E, Molina-Hernández A, González-Barrios FJ. A rapid spread of the stony coral tissue loss disease outbreak in the Mexican Caribbean. PeerJ. 2019;7: e8069. doi:10.7717/peerj.8069

26. Weil E, Hernández-Delgado E, Gonzalez M, Williams S, Suleimán S, Figuerola M, et al. Spread of the new coral disease “SCTLD” into the Caribbean: implications for Puerto Rico. Reef Encounter. 201934.

27. Brandt ME, Ennis RS, Meiling SS, Townsend J, Cobleigh K, Glahn A, et al. The Emergence and Initial Impact of Stony Coral Tissue Loss Disease (SCTLD) in the United States Virgin Islands. Front Mar Sci. 2021;8. Available: https://www.frontiersin.org/articles/10.3389/fmars.2021.715329

28. Evans JS, Paul VJ, Ushijima B, Pitts KA, Kellogg CA. Investigating microbial size classes associated with the transmission of stony coral tissue loss disease (SCTLD). PeerJ. 2023;11: e15836. doi:10.7717/peerj.15836

29. Hernandez-Agreda A, Gates RD, Ainsworth TD. Defining the Core Microbiome in Corals’ Microbial Soup. Trends Microbiol. 2017;25: 125–140. doi:10.1016/j.tim.2016.11.003

30. Egan S, Gardiner M. Microbial Dysbiosis: Rethinking Disease in Marine Ecosystems. Front Microbiol. 2016;7. doi:10.3389/fmicb.2016.00991

31. Rosales SM, Clark AS, Huebner LK, Ruzicka RR, Muller EM. Rhodobacterales and Rhizobiales Are Associated With Stony Coral Tissue Loss Disease and Its Suspected Sources of Transmission. Front Microbiol. 2020;11. Available: https://www.frontiersin.org/articles/10.3389/fmicb.2020.00681

32. Becker CC, Brandt M, Miller CA, Apprill A. Microbial bioindicators of Stony Coral Tissue Loss Disease identified in corals and overlying waters using a rapid field-based sequencing approach. Environ Microbiol. 2022;24: 1166–1182. doi:10.1111/1462-2920.15718

33. Huntley N, Brandt ME, Becker CC, Miller CA, Meiling SS, Correa AMS, et al. Experimental transmission of Stony Coral Tissue Loss Disease results in differential microbial responses within coral mucus and tissue. ISME Commun. 2022;2: 1–11. doi:10.1038/s43705-022-00126-3

34. Meyer JL, Castellanos-Gell J, Aeby GS, Häse CC, Ushijima B, Paul VJ. Microbial Community Shifts Associated With the Ongoing Stony Coral Tissue Loss Disease Outbreak on the Florida Reef Tract. Front Microbiol. 2019;10: 2244. doi:10.3389/fmicb.2019.02244

35. Ushijima B, Meyer JL, Thompson S, Pitts K, Marusich MF, Tittl J, et al. Disease Diagnostics and Potential Coinfections by Vibrio coralliilyticus During an Ongoing Coral Disease Outbreak in Florida. Front Microbiol. 2020;11. Available: https://www.frontiersin.org/articles/10.3389/fmicb.2020.569354

36. Meiling SS, Muller EM, Lasseigne D, Rossin A, Veglia AJ, MacKnight N, et al. Variable Species Responses to Experimental Stony Coral Tissue Loss Disease (SCTLD) Exposure. Front Mar Sci. 2021;8. Available: https://www.frontiersin.org/article/10.3389/fmars.2021.670829

37. Rosales SM, Huebner LK, Clark AS, McMinds R, Ruzicka RR, Muller EM. Bacterial Metabolic Potential and Micro-Eukaryotes Enriched in Stony Coral Tissue Loss Disease Lesions. Front Mar Sci. 2022;8. Available: https://www.frontiersin.org/articles/10.3389/fmars.2021.776859

38. Studivan MS, Rossin AM, Rubin E, Soderberg N, Holstein DM, Enochs IC. Reef Sediments Can Act As a Stony Coral Tissue Loss Disease Vector. Front Mar Sci. 2022;8. Available: https://www.frontiersin.org/articles/10.3389/fmars.2021.815698

39. Work TM, Weatherby TM, Landsberg JH, Kiryu Y, Cook SM, Peters EC. Viral-Like Particles Are Associated With Endosymbiont Pathology in Florida Corals Affected by Stony Coral Tissue Loss Disease. Front Mar Sci. 2021;8: 750658. doi:10.3389/fmars.2021.750658

40. Veglia AJ, Beavers KM, Van Buren EW, Meiling SS, Muller EM, Smith TB, et al. Alphaflexivirus Genomes in Stony Coral Tissue Loss Disease-Affected, Disease-Exposed, and Disease-Unexposed Coral Colonies in the U.S. Virgin Islands. Microbiol Resour Announc. 2022;11: e01199–21. doi:10.1128/mra.01199-21

41. Howe-Kerr LI, Knochel AM, Meyer MD, Sims JA, Karrick CE, Grupstra CGB, et al. Filamentous virus-like particles are present in coral dinoflagellates across genera and ocean basins. ISME J. 2023;17: 2389–2402. doi:10.1038/s41396-023-01526-6

42. Forrester GE, Arton L, Horton A, Nickles K, Forrester LM. Antibiotic Treatment Ameliorates the Impact of Stony Coral Tissue Loss Disease (SCTLD) on Coral Communities. Front Mar Sci. 2022;9. Available: https://www.frontiersin.org/articles/10.3389/fmars.2022.859740

43. Neely KL, Macaulay KA, Hower EK, Dobler MA. Effectiveness of topical antibiotics in treating corals affected by Stony Coral Tissue Loss Disease. PeerJ. 2020;8. doi:10.7717/peerj.9289

44. Shilling EN, Combs IR, Voss JD. Assessing the effectiveness of two intervention methods for stony coral tissue loss disease on Montastraea cavernosa. Sci Rep. 2021;11: 8566. doi:10.1038/s41598-021-86926-4

45. Walker BK, Turner NR, Noren HKG, Buckley SF, Pitts KA. Optimizing Stony Coral Tissue Loss Disease (SCTLD) Intervention Treatments on Montastraea cavernosa in an Endemic Zone. Front Mar Sci. 2021;8. Available: https://www.frontiersin.org/articles/10.3389/fmars.2021.666224

46. Landsberg JH, Kiryu Y, Peters EC, Wilson PW, Perry N, Waters Y, et al. Stony Coral Tissue Loss Disease in Florida Is Associated With Disruption of Host–Zooxanthellae Physiology. Front Mar Sci. 2020;7: 576013. doi:10.3389/fmars.2020.576013

47. Deutsch JM, Jaiyesimi OA, Pitts KA, Houk J, Ushijima B, Walker BK, et al. Metabolomics of Healthy and Stony Coral Tissue Loss Disease Affected Montastraea cavernosa Corals. Front Mar Sci. 2021;8. Available: https://www.frontiersin.org/articles/10.3389/fmars.2021.714778

48. Beavers KM, Van Buren EW, Rossin AM, Emery MA, Veglia AJ, Karrick CE, et al. Stony coral tissue loss disease induces transcriptional signatures of in situ degradation of dysfunctional Symbiodiniaceae. Nat Commun. 2023;14: 2915. doi:10.1038/s41467-023-38612-4

49. Traylor-Knowles N, Connelly MT, Young BD, Eaton K, Muller EM, Paul VJ, et al. Gene Expression Response to Stony Coral Tissue Loss Disease Transmission in M. cavernosa and O. faveolata From Florida. Front Mar Sci. 2021;8: 681563. doi:10.3389/fmars.2021.681563

50. Studivan MS, Eckert RJ, Shilling E, Soderberg N, Enochs IC, Voss JD. Stony coral tissue loss disease intervention with amoxicillin leads to a reversal of disease-modulated gene expression pathways. Mol Ecol. 2023;32: 5394–5413. doi:10.1111/mec.17110

51. Gliddon HD, Herberg JA, Levin M, Kaforou M. Genome-wide host RNA signatures of infectious diseases: discovery and clinical translation. Immunology. 2018;153: 171–178. doi:10.1111/imm.12841

52. Das P, Roychowdhury A, Das S, Roychoudhury S, Tripathy S. sigFeature: Novel Significant Feature Selection Method for Classification of Gene Expression Data Using Support Vector Machine and t Statistic. Front Genet. 2020;11. Available: https://www.frontiersin.org/articles/10.3389/fgene.2020.00247

53. Burzykowski T, Rousseau A-J, Geubbelmans M, Valkenborg D. Introduction to machine learning. Am J Orthod Dentofacial Orthop. 2023;163: 732–734. doi:10.1016/j.ajodo.2023.02.005

54. Bellman R, Kalaba R. On adaptive control processes. IRE Trans Autom Control. 1959;4: 1–9. doi:10.1109/TAC.1959.1104847

55. Guyon I, Weston J, Barnhill S, Vapnik V. Gene Selection for Cancer Classification using Support Vector Machines. Mach Learn. 2002;46: 389–422. doi:10.1023/A:1012487302797

56. Liang Y, Zhang F, Wang J, Joshi T, Wang Y, Xu D. Prediction of Drought-Resistant Genes in Arabidopsis thaliana Using SVM-RFE. PLOS ONE. 2011;6: e21750. doi:10.1371/journal.pone.0021750

57. Richhariya B, Tanveer M, Rashid AH. Diagnosis of Alzheimer’s disease using universum support vector machine based recursive feature elimination (USVM-RFE). Biomed Signal Process Control. 2020;59: 101903. doi:10.1016/j.bspc.2020.101903

58. Davies SW, Ries JB, Marchetti A, Castillo KD. Symbiodinium Functional Diversity in the Coral Siderastrea siderea Is Influenced by Thermal Stress and Reef Environment, but Not Ocean Acidification. Front Mar Sci. 2018;5: 150. doi:10.3389/fmars.2018.00150

59. Simão FA, Waterhouse RM, Ioannidis P, Kriventseva EV, Zdobnov EM. BUSCO: assessing genome assembly and annotation completeness with single-copy orthologs. Bioinformatics. 2015;31: 3210–3212. doi:10.1093/bioinformatics/btv351

60. Bushnell B. BBMap: A Fast, Accurate, Splice-Aware Aligner. Lawrence Berkeley National Lab. (LBNL), Berkeley, CA (United States); 2014 Mar. Report No.: LBNL-7065E. Available: https://www.osti.gov/biblio/1241166

61. Paysan-Lafosse T, Blum M, Chuguransky S, Grego T, Pinto BL, Salazar GA, et al. InterPro in 2022. Nucleic Acids Res. 2022;51: D418–D427. doi:10.1093/nar/gkac993

62. Uddin S, Khan A, Hossain ME, Moni MA. Comparing different supervised machine learning algorithms for disease prediction. BMC Med Inform Decis Mak. 2019;19: 281. doi:10.1186/s12911-019-1004-8

63. Imbs AB, Dembitsky VM. Coral Lipids. Mar Drugs. 2023;21: 539. doi:10.3390/md21100539

64. Papina M, Meziane T, van Woesik R. Symbiotic zooxanthellae provide the host-coral Montipora digitata with polyunsaturated fatty acids. Comp Biochem Physiol B Biochem Mol Biol. 2003;135: 533–537. doi:10.1016/S1096-4959(03)00118-0

65. Clark DA, Coker R. Molecules in focus Transforming growth factor-beta (TGF-β). Int J Biochem Cell Biol. 1998;30: 293–298. doi:10.1016/S1357-2725(97)00128-3

66. Detournay O, Schnitzler CE, Poole A, Weis VM. Regulation of cnidarian–dinoflagellate mutualisms: Evidence that activation of a host TGFβ innate immune pathway promotes tolerance of the symbiont. Dev Comp Immunol. 2012;38: 525–537. doi:10.1016/j.dci.2012.08.008

67. Berthelier J, Schnitzler CE, Wood-Charlson EM, Poole AZ, Weis VM, Detournay O. Implication of the host TGFβ pathway in the onset of symbiosis between larvae of the coral Fungia scutaria and the dinoflagellate Symbiodinium sp. (clade C1f). Coral Reefs. 2017;36: 1263–1268. doi:10.1007/s00338-017-1621-6

68. Fuess LE, Mann WT, Jinks LR, Brinkhuis V, Mydlarz LD. Transcriptional analyses provide new insight into the late-stage immune response of a diseased Caribbean coral. R Soc Open Sci. 2018;5: 172062. doi:10.1098/rsos.172062

69. Rosenstiel P, Sina C, End C, Renner M, Lyer S, Till A, et al. Regulation of *DMBT1* via NOD2 and TLR4 in Intestinal Epithelial Cells Modulates Bacterial Recognition and Invasion. J Immunol. 2007;178: 8203–8211. doi:10.4049/jimmunol.178.12.8203

70. McDowell IC, Nikapitiya C, Aguiar D, Lane CE, Istrail S, Gomez-Chiarri M. Transcriptome of American Oysters, Crassostrea virginica, in Response to Bacterial Challenge: Insights into Potential Mechanisms of Disease Resistance. PLOS ONE. 2014;9: e105097. doi:10.1371/journal.pone.0105097

71. Fuess LE, Pinzόn C. JH, Weil E, Mydlarz LD. Associations between transcriptional changes and protein phenotypes provide insights into immune regulation in corals. Dev Comp Immunol. 2016;62: 17–28. doi:10.1016/j.dci.2016.04.017

72. Riesgo A, Peterson K, Richardson C, Heist T, Strehlow B, McCauley M, et al. Transcriptomic analysis of differential host gene expression upon uptake of symbionts: a case study with Symbiodinium and the major bioeroding sponge Cliona varians. BMC Genomics. 2014;15: 376. doi:10.1186/1471-2164-15-376

73. Wright RM, Kenkel CD, Dunn CE, Shilling EN, Bay LK, Matz MV. Intraspecific differences in molecular stress responses and coral pathobiome contribute to mortality under bacterial challenge in Acropora millepora. Sci Rep. 2017;7: 2609. doi:10.1038/s41598-017-02685-1

74. Tsugeki R, Ditengou FA, Sumi Y, Teale W, Palme K, Okada K. NO VEIN Mediates Auxin-Dependent Specification and Patterning in the Arabidopsis Embryo, Shoot, and Root. Plant Cell. 2009;21: 3133–3151. doi:10.1105/tpc.109.068841

75. Gonen N, Meller A, Sabath N, Shalgi R. Amino Acid Biosynthesis Regulation during Endoplasmic Reticulum Stress Is Coupled to Protein Expression Demands. iScience. 2019;19: 204–213. doi:10.1016/j.isci.2019.07.022

76. Howe R, Kelly M, Jimah J, Hodge D, Odom AR. Isoprenoid Biosynthesis Inhibition Disrupts Rab5 Localization and Food Vacuolar Integrity in Plasmodium falciparum. Eukaryot Cell. 2013;12: 215–223. doi:10.1128/ec.00073-12

77. Fransolet D, Roberty S, Plumier J-C. Establishment of endosymbiosis: The case of cnidarians and Symbiodinium. J Exp Mar Biol Ecol. 2012;420–421: 1–7. doi:10.1016/j.jembe.2012.03.015

78. Chen MC, Cheng YM, Hong MC, Fang LS. Molecular cloning of Rab5 (ApRab5) in Aiptasia pulchella and its retention in phagosomes harboring live zooxanthellae. Biochem Biophys Res Commun. 2004;324: 1024–1033. 10.1016/j.bbrc.2004.09.151

79. Hashim S, Mukherjee K, Raje M, Basu SK, Mukhopadhyay A. Live Salmonella Modulate Expression of Rab Proteins to Persist in a Specialized Compartment and Escape Transport to Lysosomes*. J Biol Chem. 2000;275: 16281–16288. doi:10.1074/jbc.275.21.16281

80. Via LE, Deretic D, Ulmer RJ, Hibler NS, Huber LA, Deretic V. Arrest of Mycobacterial Phagosome Maturation Is Caused by a Block in Vesicle Fusion between Stages Controlled by rab5 and rab7*. J Biol Chem. 1997;272: 13326–13331. doi:10.1074/jbc.272.20.13326

81. Alvarez-Dominguez C, Barbieri AM, Berón W, Wandinger-Ness A, Stahl PD. Phagocytosed Live Listeria monocytogenes Influences Rab5-regulated in Vitro Phagosome-Endosome Fusion*. J Biol Chem. 1996;271: 13834–13843. doi:10.1074/jbc.271.23.13834

82. Brown B, Bythell J. Perspectives on mucus secretion in reef corals. Mar Ecol Prog Ser. 2005;296: 291–309. doi:10.3354/meps296291

83. Lamberti G, Gügel IL, Meurer J, Soll J, Schwenkert S. The Cytosolic Kinases STY8, STY17, and STY46 Are Involved in Chloroplast Differentiation in Arabidopsis1[W]. Plant Physiol. 2011;157: 70–85. doi:10.1104/pp.111.182774

84. Emery MA, Dimos BA, Mydlarz LD. Cnidarian Pattern Recognition Receptor Repertoires Reflect Both Phylogeny and Life History Traits. Front Immunol. 2021;12. Available: https://www.frontiersin.org/articles/10.3389/fimmu.2021.689463

85. Brown GD, Willment JA, Whitehead L. C-type lectins in immunity and homeostasis. Nat Rev Immunol. 2018;18: 374–389. doi:10.1038/s41577-018-0004-8

86. Davy SK, Allemand D, Weis VM. Cell Biology of Cnidarian-Dinoflagellate Symbiosis. Microbiol Mol Biol Rev. 2012;76. doi:10.1128/mmbr.05014-11

87. Mansfield KM, Carter NM, Nguyen L, Cleves PA, Alshanbayeva A, Williams LM, et al. Transcription factor NF-κB is modulated by symbiotic status in a sea anemone model of cnidarian bleaching. Sci Rep. 2017;7: 16025. doi:10.1038/s41598-017-16168-w

88. Rivera HE, Davies SW. Symbiosis maintenance in the facultative coral, Oculina arbuscula, relies on nitrogen cycling, cell cycle modulation, and immunity. Sci Rep. 2021;11: 21226. doi:10.1038/s41598-021-00697-6

89. Weis VM. Cell Biology of Coral Symbiosis: Foundational Study Can Inform Solutions to the Coral Reef Crisis. Integr Comp Biol. 2019;59: 845–855. doi:10.1093/icb/icz067

90. Mansfield KM, Gilmore TD. Innate immunity and cnidarian-Symbiodiniaceae mutualism. Dev Comp Immunol. 2019;90: 199–209. doi:10.1016/j.dci.2018.09.020

91. Emery MA, Beavers KM, Van Buren EW, Batiste R, Dimos B, Pellegrino MW, et al. Trade-off between photosymbiosis and innate immunity influences cnidarian’s response to pathogenic bacteria. Proc R Soc B Biol Sci. 2024;291: 20240428. doi:10.1098/rspb.2024.0428

92. Zhao YG, Zhang H. Autophagosome maturation: An epic journey from the ER to lysosomes. J Cell Biol. 2018;218: 757–770. doi:10.1083/jcb.201810099

93. Downs CA, Kramarsky-Winter E, Martinez J, Kushmaro A, Woodley CM, Loya Y, et al. Symbiophagy as a cellular mechanism for coral bleaching. Autophagy. 2009;5: 211–216. doi:10.4161/auto.5.2.7405

94. Desalvo MK, Voolstra CR, Sunagawa S, Schwarz JA, Stillman JH, Coffroth MA, et al. Differential gene expression during thermal stress and bleaching in the Caribbean coral Montastraea faveolata. Mol Ecol. 2008;17: 3952–3971. doi:10.1111/j.1365-294X.2008.03879.x

95. Smith EG, D’Angelo C, Salih A, Wiedenmann J. Screening by coral green fluorescent protein (GFP)-like chromoproteins supports a role in photoprotection of zooxanthellae. Coral Reefs. 2013;32: 463–474. doi:10.1007/s00338-012-0994-9

96. Barshis DJ, Ladner JT, Oliver TA, Seneca FO, Traylor-Knowles N, Palumbi SR. Genomic basis for coral resilience to climate change. Proc Natl Acad Sci. 2013;110: 1387–1392. doi:10.1073/pnas.1210224110

97. Lesser MP. Oxidative stress causes coral bleaching during exposure to elevated temperatures. Coral Reefs. 1997;16: 187–192. doi:https://doi-org.ezproxy.uta.edu/10.1007/s003380050073

98. Weis VM. Cellular mechanisms of Cnidarian bleaching: stress causes the collapse of symbiosis. J Exp Biol. 2008;211: 3059–3066. doi:10.1242/jeb.009597

99. Ray P, Reddy SS, Banerjee T. Various dimension reduction techniques for high dimensional data analysis: a review. Artif Intell Rev. 2021;54: 3473–3515. doi:10.1007/s10462-020-09928-0

100. Yousef M, Kumar A, Bakir-Gungor B. Application of Biological Domain Knowledge Based Feature Selection on Gene Expression Data. Entropy. 2021;23: 2. doi:10.3390/e23010002

101. Ambroise C, McLachlan GJ. Selection bias in gene extraction on the basis of microarray gene-expression data. Proc Natl Acad Sci. 2002;99: 6562–6566. doi:10.1073/pnas.102102699

102. Yates LA, Aandahl Z, Richards SA, Brook BW. Cross validation for model selection: A review with examples from ecology. Ecol Monogr. 2023;93: e1557. doi:10.1002/ecm.1557

103. SCTLD Case Definition. Florida Coral Disease Response Research & Epidemiology Team; 2018. Available: https://floridadep.gov/sites/default/files/Copy%20of%20StonyCoralTissueLossDisease_CaseDefinition%20final%2010022018.pdf

104. Stanzione D, West J, Evans RT, Minyard T, Ghattas O, Panda DK. Frontera: The Evolution of Leadership Computing at the National Science Foundation. Practice and Experience in Advanced Research Computing. New York, NY, USA: Association for Computing Machinery; 2020. pp. 106–111. doi:10.1145/3311790.3396656

105. Chen S, Zhou Y, Chen Y, Gu J. fastp: an ultra-fast all-in-one FASTQ preprocessor. Bioinformatics. 2018;34: i884–i890. doi:10.1093/bioinformatics/bty560

106. Haas BJ, Papanicolaou A, Yassour M, Grabherr M, Blood PD, Bowden J, et al. De novo transcript sequence reconstruction from RNA-seq using the Trinity platform for reference generation and analysis. Nat Protoc. 2013;8: 1494–1512. doi:10.1038/nprot.2013.084

107. Davies SW, Marchetti A, Ries JB, Castillo KD. Thermal and pCO2 Stress Elicit Divergent Transcriptomic Responses in a Resilient Coral. Front Mar Sci. 2016;3. Available: https://www.frontiersin.org/article/10.3389/fmars.2016.00112

108. Beavers KM. Master Coral database used in USVI SCTLD Transmission Experiment Gene Expression Analysis. Zenodo; 2023. doi:10.5281/zenodo.7838980

109. Sayers EW, Bolton EE, Brister JR, Canese K, Chan J, Comeau DC, et al. Database resources of the National Center for Biotechnology Information. Nucleic Acids Res. 2021;50: D20–D26. doi:10.1093/nar/gkab1112

110. Haas, BJ. TransDecoder. In: TransDecoder [Internet]. 5 May [cited 24 Aug 2023]. Available: https://github.com/TransDecoder/TransDecoder

111. Huang Y, Niu B, Gao Y, Fu L, Li W. CD-HIT Suite: a web server for clustering and comparing biological sequences. Bioinformatics. 2010;26: 680–682. doi:10.1093/bioinformatics/btq003

112. Bayer T, Aranda M, Sunagawa S, Yum LK, DeSalvo MK, Lindquist E, et al. Symbiodinium Transcriptomes: Genome Insights into the Dinoflagellate Symbionts of Reef-Building Corals. Moustafa A, editor. PLoS ONE. 2012;7: e35269. doi:10.1371/journal.pone.0035269

113. Parkinson JE, Baumgarten S, Michell CT, Baums IB, LaJeunesse TC, Voolstra CR. Gene Expression Variation Resolves Species and Individual Strains among Coral-Associated Dinoflagellates within the Genus *Symbiodinium*. Genome Biol Evol. 2016;8: 665–680. doi:10.1093/gbe/evw019

114. Bellantuono AJ, Dougan KE, Granados-Cifuentes C, Rodriguez-Lanetty M. Free-living and symbiotic lifestyles of a thermotolerant coral endosymbiont display profoundly distinct transcriptomes under both stable and heat stress conditions. Mol Ecol. 2019;28: 5265–5281. doi:10.1111/mec.15300

115. Patro R, Duggal G, Love MI, Irizarry RA, Kingsford C. Salmon provides fast and bias-aware quantification of transcript expression. Nat Methods. 2017;14: 417–419. doi:10.1038/nmeth.4197

116. R Core Team. R: A language and environment for statistical computing. R Foundation for Statistical Computing, Vienna, Austria; 2022. Available: https://www.R-project.org/

117. Soneson C, Love MI, Robinson MD. Differential analyses for RNA-seq: transcript-level estimates improve gene-level inferences. F1000Research. 2016;4: 1521. doi:10.12688/f1000research.7563.2

118. Love MI, Huber W, Anders S. Moderated estimation of fold change and dispersion for RNA-seq data with DESeq2. Genome Biol. 2014;15: 550. doi:10.1186/s13059-014-0550-8

119. Szklarczyk D, Gable AL, Lyon D, Junge A, Wyder S, Huerta-Cepas J, et al. STRING v11: protein-protein association networks with increased coverage, supporting functional discovery in genome-wide experimental datasets. Nucleic Acids Res. 2019;47: D607–D613. doi:10.1093/nar/gky1131

