## Supplementary figures and images for "Characterizing gene expression profiles of various tissue states in stony coral tissue loss disease using a feature selection algorithm"

### Fig S1

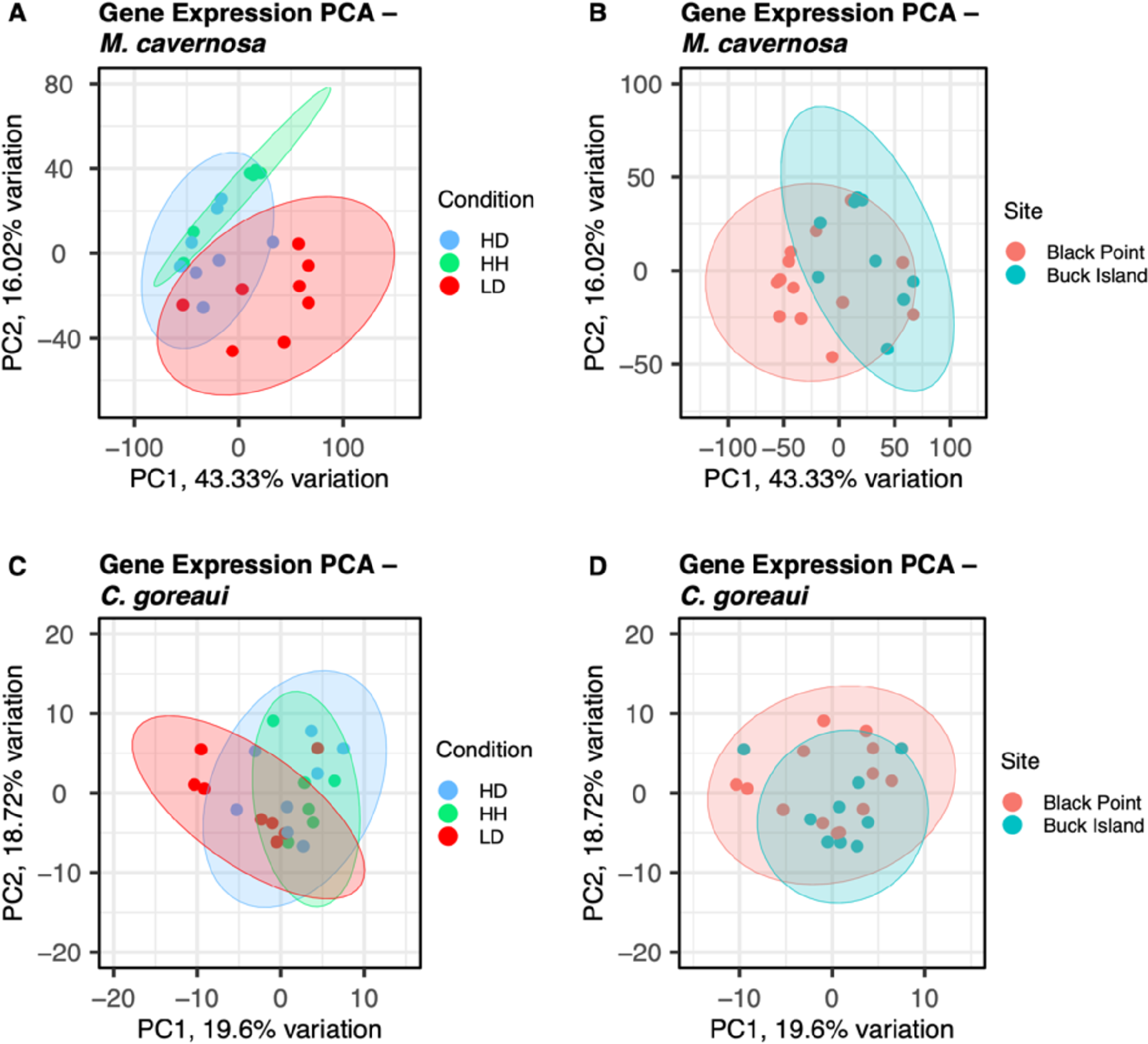

### Fig S2

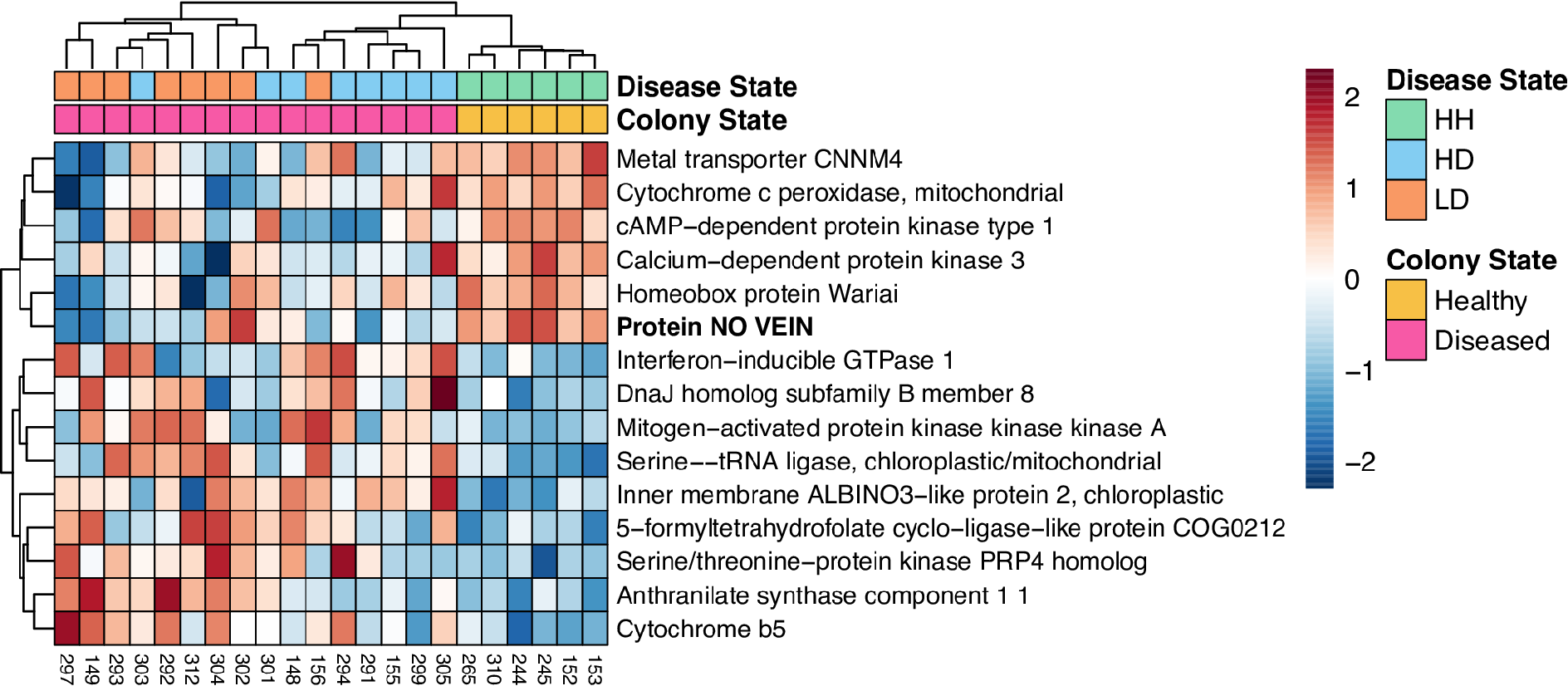

### Fig S3

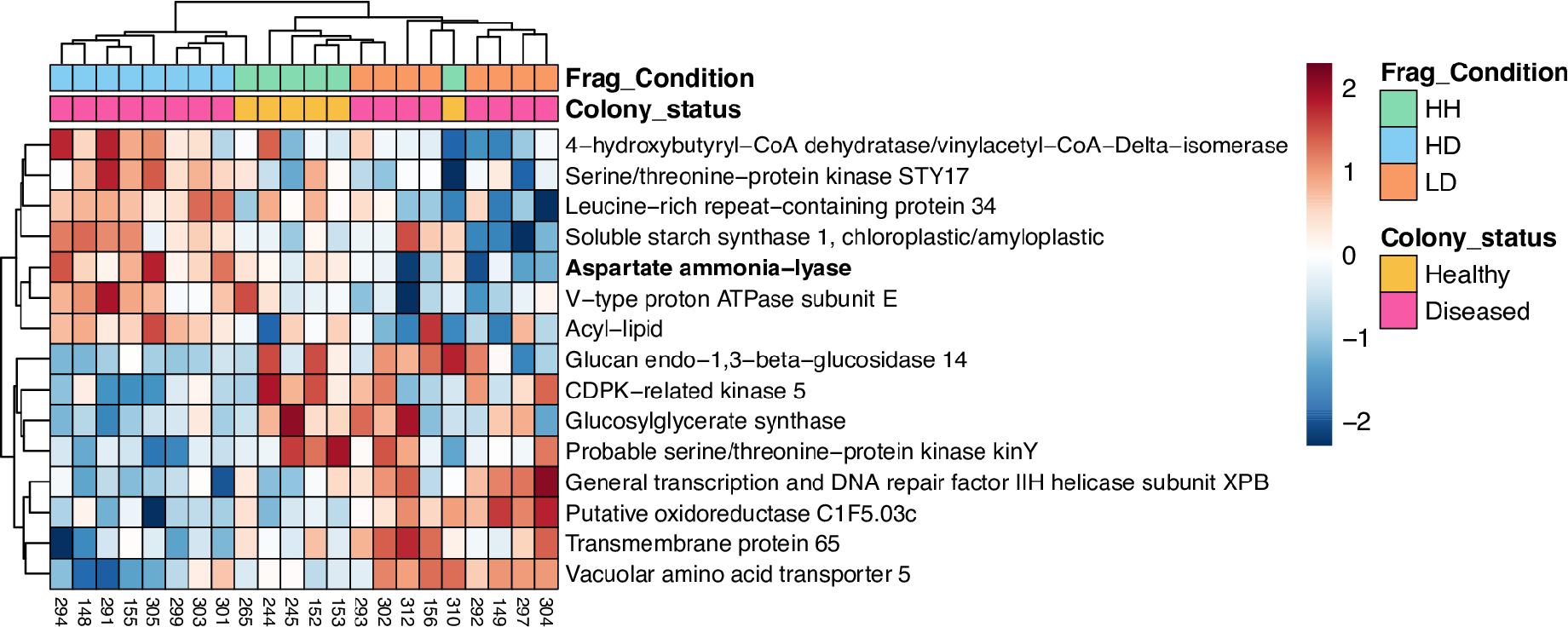

### Fig S4

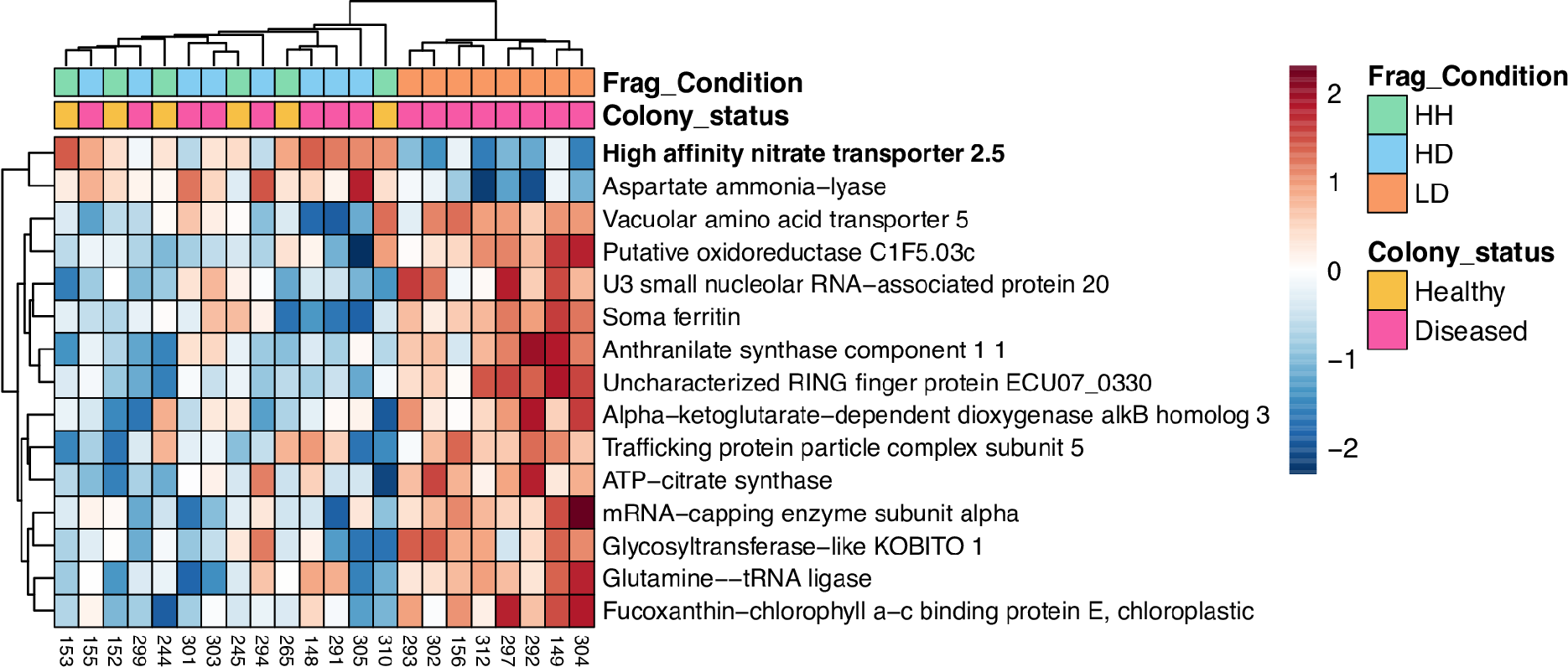
